# Inhalable textile microplastic fibers impair lung repair

**DOI:** 10.1101/2021.01.25.428144

**Authors:** F. van Dijk, S. Song, G.W.A van Eck, X. Wu, I.S.T. Bos, D.H.A. Boom, I.M. Kooter, D.C.J. Spierings, R. Wardenaar, M. Cole, A. Salvati, R. Gosens, B.N. Melgert

**Author notes:** Corresponding author: Prof Dr Barbro N. Melgert, Groningen Research Institute of Pharmacy, Department of Molecular Pharmacology, University of Groningen, A. Deusinglaan 1, 9713 AV Groningen, the Netherlands.

## Abstract

Synthetic textiles shed fibers that accumulate indoors and this results in continuous exposure when indoors. High exposure to microplastic fibers in nylon flock workers has been linked to the development of airway and interstitial lung disease, but the exact health effects of microplastic fibers on the lungs are unknown. Here we determined effects of polyester and nylon textile microplastic fibers on airway and alveolar epithelial cells using human and murine lung organoids. We observed that particularly nylon microfibers had a negative impact on the growth and development of airway organoids. We demonstrated that this effect was mediated by components leaking from nylon. Moreover, our data suggested that microplastic textile fibers may especially harm the developing airways or airways undergoing repair. Our results call for a need to assess exposure and inhalation levels in indoor environments to accurately determine the actual risk of these fibers to human health.

**Teaser:** Airborne fibers shed from synthetic textiles, in particular nylon, can inhibit repair of the cells coating the airways

## Introduction

Plastic pollution is a pressing global concern and microplastics are a significant part of this problem (*1*). High amounts of microplastics have been found in marine environments, air, soils, plants, and animals, which illustrates how omnipresent this relatively recent pollution actually is (*2*). Microplastic pollution derives from personal care products, synthetic clothes, and degradation of macroplastics (*3, 4*). Synthetic textile fibers are one of the most prevalent types of microplastic waste observed, with an annual production of 60 million metric tons, which equals 16% of the world’s plastic pollution (*1*). These fibers are typically composed of nylon or polyester and are released into the environment by wear and tear and during washing and drying of garments (*4–6*).

The ubiquitous nature of microplastics in the environment inevitably leads to human exposure, which can occur through two main routes (*7, 8*). Firstly, through ingestion of contaminated food and water and secondly via inhalation. Microplastics have been reported both in indoor and outdoor air, with levels indoors being 2-5 times higher as compared to outdoors (*9, 10*). Whether or not microfibers can deposit in lung tissue largely depends on the aerodynamic diameter of the fibers (*7, 11*). Lung deposition is most efficiently achieved with aerodynamic diameters between 1-10 μm (*12*), however, these sizes are difficult to quantify in environmental samples due to limitations of the analytical techniques (*9, 13*). Yet, plastic microfibers have been found in human lung tissue, suggesting inhalation does indeed take place (*14*). Furthermore, several studies from workers in synthetic textile, flock and (poly)vinyl chloride industries suggest that inhalation of such microfibers is harmful, as around 30% of factory workers developed work-related airway and interstitial lung disease (*15–23*). Moreover, exposure to particulate matter in air pollution, also containing microplastics (*24–26*), has been associated with higher risk of developing asthma and an increase in asthma symptoms in areas with higher levels of particulate matter air pollution (*27–29*).

Despite the potential capacity for microplastic fibers to contribute to respiratory diseases, the health effects are greatly understudied and information providing evidence of potential human health effects of inhaled microplastics is lacking (*9, 30, 31*). In the present study, we therefore explored whether textile microplastic fibers can cause damage to lung tissue. As epithelial cells are the first to come into contact with inhaled fibers, we investigated effects of polyester and nylon microfibers on lung epithelial proliferation, differentiation, and repair processes. For this we used lung organoids that are grown from primary lung epithelial progenitor cells with support of a lung fibroblast cell line (*32, 33*). The epithelial progenitors can develop into organoids consisting of alveolar epithelial cells or organoids consisting of airway epithelial cells with help of growth factors produced by fibroblasts. We found that in particular nylon microfibers negatively impacted developing airway organoids, while developing alveolar organoids and already developed organoids of both types appeared to be less affected. This negative effect was caused by still unknown leachates from nylon that particularly inhibit differentiation of airway epithelial cells. We therefore call for assessment of exposure levels in indoor environments and actual lung deposition to accurately determine the risk of these fibers to human health.

## Results

### Characterization of reference microfibers

To produce reference textile fibers that resemble microplastics found in our indoor environments, we used a method previous described by us to reproducibly generate fibers of specific lengths (*34*). We particularly focused on polyester and nylon, because these are the most abundant types of microplastics indoors (*9, 35–38*). As we spend the majority of our time indoors, we may therefore be exposed most to these types of microplastics (*39*). Fibers are commonly defined as having a length to diameter ratio of 3:1 (*40*). The fibers we produced had a median size of 15×52 µm for polyester and 12×31 µm for nylon (Table S1). Using scanning electron microscopy (SEM) we found these fibers to be rod-shaped with a circular cross-section and had a smooth surface (Figure 1A and B). For our polyester fibers energy dispersive X-ray (EDX) analysis confirmed the presence of carbon and oxygen (Figure S1A). The recorded micro-Fourier transform infrared (μFTIR) spectrum showed characteristic absorbance peaks of polyester (Figure S1B). The EDX spectrum for nylon confirmed the presence of carbon, nitrogen and oxygen (Figure S1C) and the μFTIR spectrum showed characteristic nylon absorbance peaks (Figure S1D).

**Figure 1.**
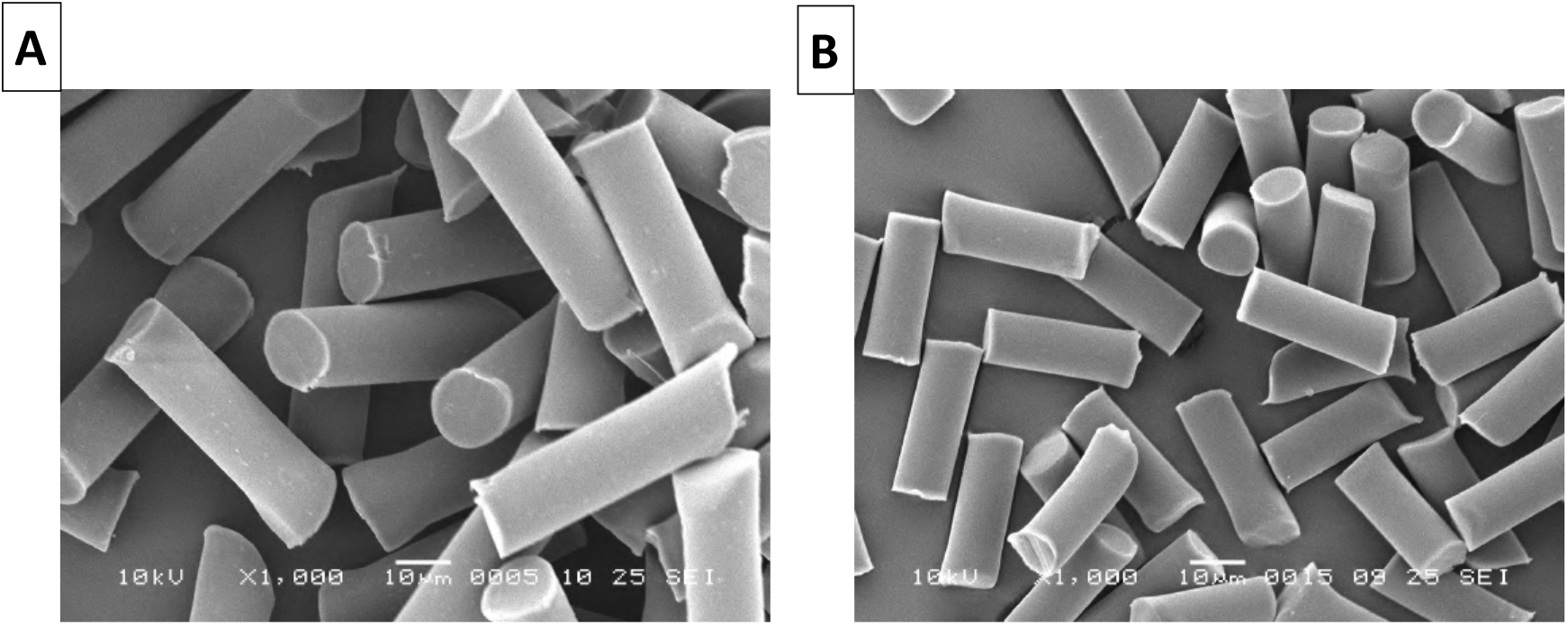
Morphology of reference microplastic fibers of standardized dimensions. Representative SEM micrographs of (**A**) polyester microfibers (15×52 µm) and (**B**) nylon microfibers (12×31 µm).

### Nylon microfibers inhibited growth of murine and human lung organoids

Possible effects of microplastic fibers on lung epithelial proliferation, differentiation, and repair processes were assessed *in vitro* using both murine and a human lung organoids. Airways are lined with ciliated pseudostratified epithelium consisting of basal cells, ciliated cells, and secretory cells like goblet cells and club cells, while alveoli consist of alveolar epithelial cells type I and II (AECI and AECII). Basal cells and club cells have stem cell-like abilities and basal cells can give rise to all important epithelial cells lining the airways, while club cells can develop into ciliated cells, goblet cells, and AECII (*41, 42*). AECII can behave as alveolar stem cells and can proliferate and develop into AECI (*43*). The lung organoids we used in these studies self-assemble from the lung epithelial progenitor/stem cells isolated from adult lung tissue, i.e. basal cells, club cells and AECII. The growth of organoids from these progenitor cells is supported by proliferation and differentiation enhancers produced by epithelial cells themselves and fibroblasts also present in our 3D cultures (*43*).

We first assessed the effects of several doses of fibers on organoid growth ranging from 2000-5000 fibers per well (Figure S2). The fibers were dispersed into liquid Matrigel at the same time as the isolated epithelial cells and fibroblasts were added, after which the Matrigel solidifies and epithelial progenitors start to form organoids. Based on these results we continued with 5000 polyester or 5000 nylon fibers per well, equivalent to 122 μg/ml polyester and 39 μg/ml nylon, as this concentration had clear effects and was on the lower end of the spectrum of concentrations used in other studies (*7*).

Murine lung organoids develop into two distinct phenotypes, i.e. acetylated α-tubulin-positive airway organoids (Figure 2A) and prosurfactant protein C-positive alveolar organoids (Figure 2B). We assessed the effects of the two types of fibers on these two structures separately. Exposure during 14 days to either polyester or nylon microfibers resulted in significantly fewer organoids (Figure 2C) compared to untreated controls (Figure 2D and E). The effect of nylon on airway organoids was most profound of the two types of plastic and the two types of structures. Moreover, both airway and alveolar organoids were significantly smaller in size following nylon microfiber exposure as compared to untreated controls (Figure 2F and G).

**Figure 2.**
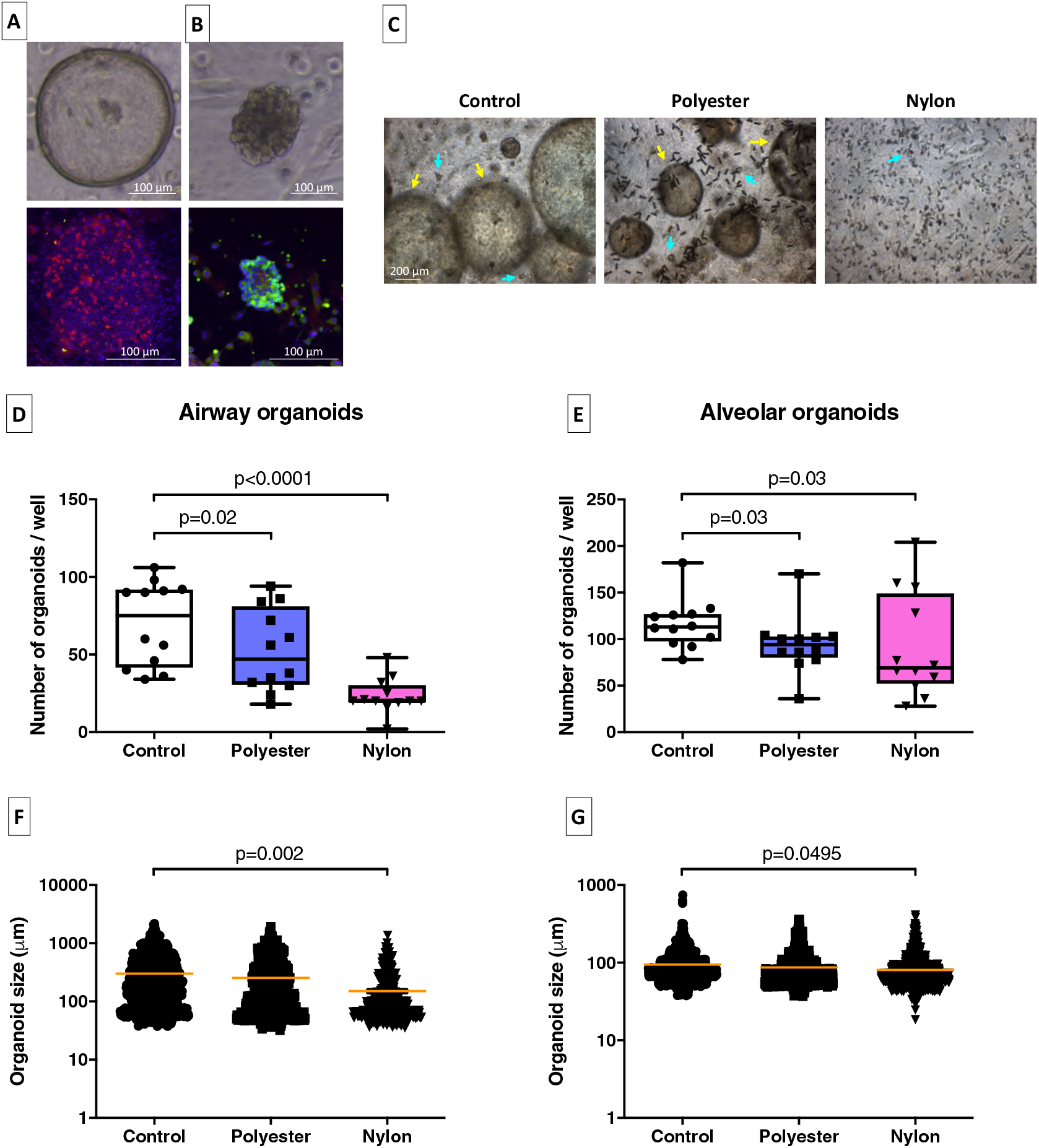
Effects of microplastic fibers on growth of murine lung organoids. Light microscopy images and fluorescence photographs of (**A**) acetylated α-tubulin-positive airway organoids (red) and (**B**) prosurfactant protein C-positive alveolar organoids (green). Nuclei were counterstained with DAPI (blue). (**C**) Representative light microscopy images of the different treatment conditions. Yellow arrows in the light microscopy images indicate airway organoids, whereas cyan arrows indicate alveolar organoids. (**D and E**) Quantification of the numbers and (**F and G**) quantification of the sizes of airway and alveolar lung organoids exposed for 14 days to no fibers, 5000 polyester, or 5000 nylon fibers (equivalent to 122 μg/ml polyester or 39 μg/ml nylon, n=12 independent isolations). Groups were compared using a Friedman test with Dunn’s correction for multiple testing. P<0.05 was considered significant.

Similar results were observed in human lung organoids, that mainly develop into alveolar organoids or mixed alveolar/airway organoids positive (Figure 3A). 14 day-exposure to nylon microfibers resulted in significantly fewer human lung organoids (Figure 3B and C), whereas the effects of polyester on organoid growth were less profound. The size of the organoids was not affected by the presence of nylon microfibers (Figure 3D).

**Figure 3.**
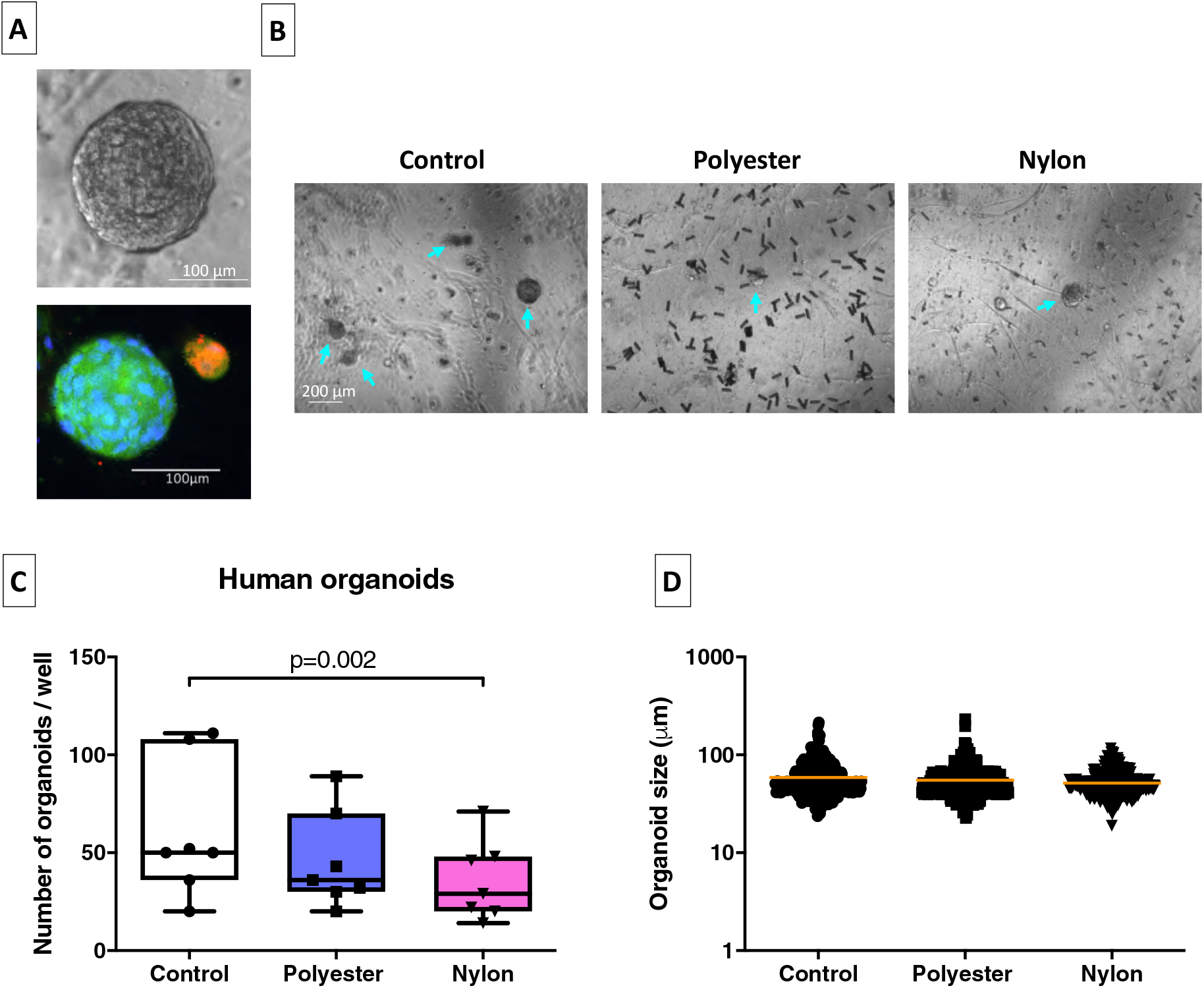
Influence of microplastic fibers on growth of human lung organoids. (**A**) The morphology of the alveolar prosurfactant protein C-positive organoids (green) and mixed acetylated α-tubulin/prosurfactant protein C-positive organoids (orange) as shown by light and fluorescence microscopy. Nuclei were counterstained with DAPI (blue). (**B**) Representative light microscopy images of all treatment conditions. Cyan arrows indicate lung organoids. (**C**) Quantification of the numbers and (**D**) sizes of human lung organoids following 14-day exposure to either no microfibers, 5000 polyester, or 5000 nylon fibers (equivalent to 122 μg/ml polyester or 39 μg/ml nylon, n=7 independent isolations). Groups were compared using a Friedman test with Dunn’s correction for multiple testing. P<0.05 was considered significant.

### Environmental microplastic fibers impaired lung organoid growth as well

Having observed these effects after exposure to reference microfibers, we next performed similar experiments using environmentally relevant polyester and nylon fibers on murine lung organoids. These were made from white polyester and nylon fabrics purchased in a local fabric store and cut to sizes approximating the reference fibers. First, we characterized morphology and chemical composition of these fibers. For polyester, we observed a more heterogeneous size distribution as compared to the reference fibers, with fibers having a median size of 17×63 µm (Figure 4A, Table S2), but a comparable EDX and µFTIR spectrum (Figure S3A and B). The nylon fibers had a disk-shaped appearance but similar dimensions as the reference nylon fibers (median 57×20 µm, Table S2). The EDX analysis revealed the expected C, N and O peaks for nylon (Figure S3C) and the µFTIR spectrum showed characteristic nylon absorbance peaks (Figure S3D).

**Figure 4.**
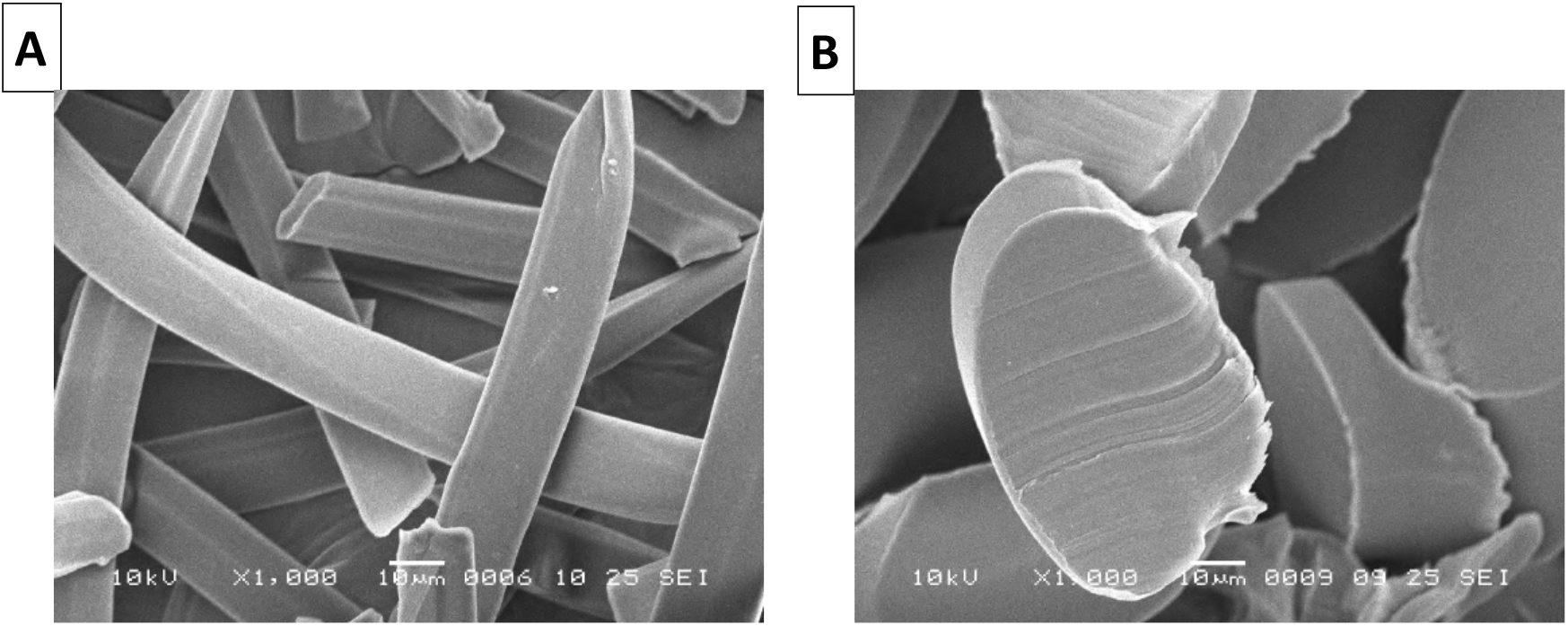
Morphology of environmental microplastic fibers. Representative SEM pictures of (**A**) polyester microfibers (17×63 µm) and (**B**) nylon microfibers (57×20 µm).

As observed with the reference fibers, exposure to environmental nylon microfibers resulted in markedly fewer lung organoids (Figure 5A, B and C) as well as smaller organoids (Figure 5D and E).

**Figure 5:**
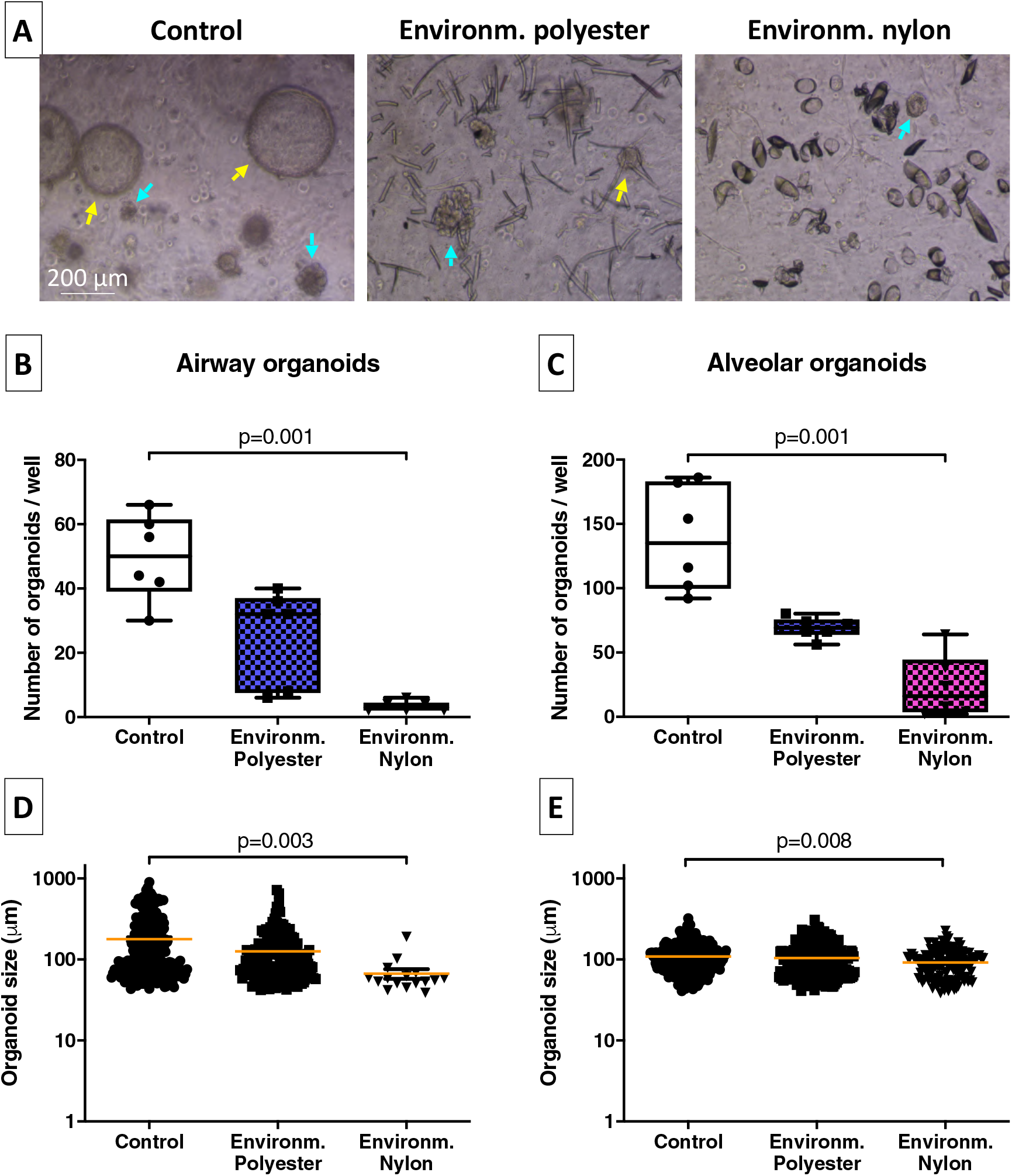
Effect of environmentally relevant textile fibers on growth of murine lung organoids. (**A**) Representative light microscopy images of all treatment conditions. Yellow arrows in the light microscopy images indicate airway organoids, whereas cyan arrows indicate alveolar organoids. (**B and C**) Quantification of the numbers and (**D and E**) sizes of airway and alveolar organoids (n=6 independent isolations) following 14-day exposure to either no microfibers, 5000 polyester or 5000 nylon microfibers (approximately equivalent to 189 μg/ml polyester or 531 μg/ml nylon). Groups were compared using a Friedman test with Dunn’s correction for multiple testing. P<0.05 was considered significant.

### Leaching nylon components caused a reduction in lung organoid growth

Since organoid growth was most affected by nylon, both reference and environmentally relevant fibers, we investigated whether this inhibition was caused by the physical presence of fibers nearby the cells or by leaching components from these nylon fibers. Therefore, we added nylon reference microfibers either on top of the Matrigel after it had set, thereby preventing direct contact with the cells, or added leachate of these fibers to the medium surrounding the Matrigel for 14 days. Interestingly, even when excluding physical contact between the fibers and the cells or simply exposing the cells to medium with leachate, the same effects on airway organoids were observed as when the forming organoids were directly exposed to the fibers. We found significantly fewer airway organoids in the presence of nylon microfibers on top of the gel or their leachate (Figure 6A and B) compared to having the fibers inside the Matrigel. The number of alveolar organoids, on the other hand, was unaffected (fibers on top) or even induced (leachate, Figure 6A and C) compared to having the fibers inside the Matrigel. Additionally, the size of these airway organoids was smaller as compared to untreated control organoids (Figure 6D), while only slightly inhibiting the size of the alveolar organoids (Figure 6E). These data suggest that specifically airway epithelial growth is inhibited by components leaching from nylon microplastics.

**Figure 6.**
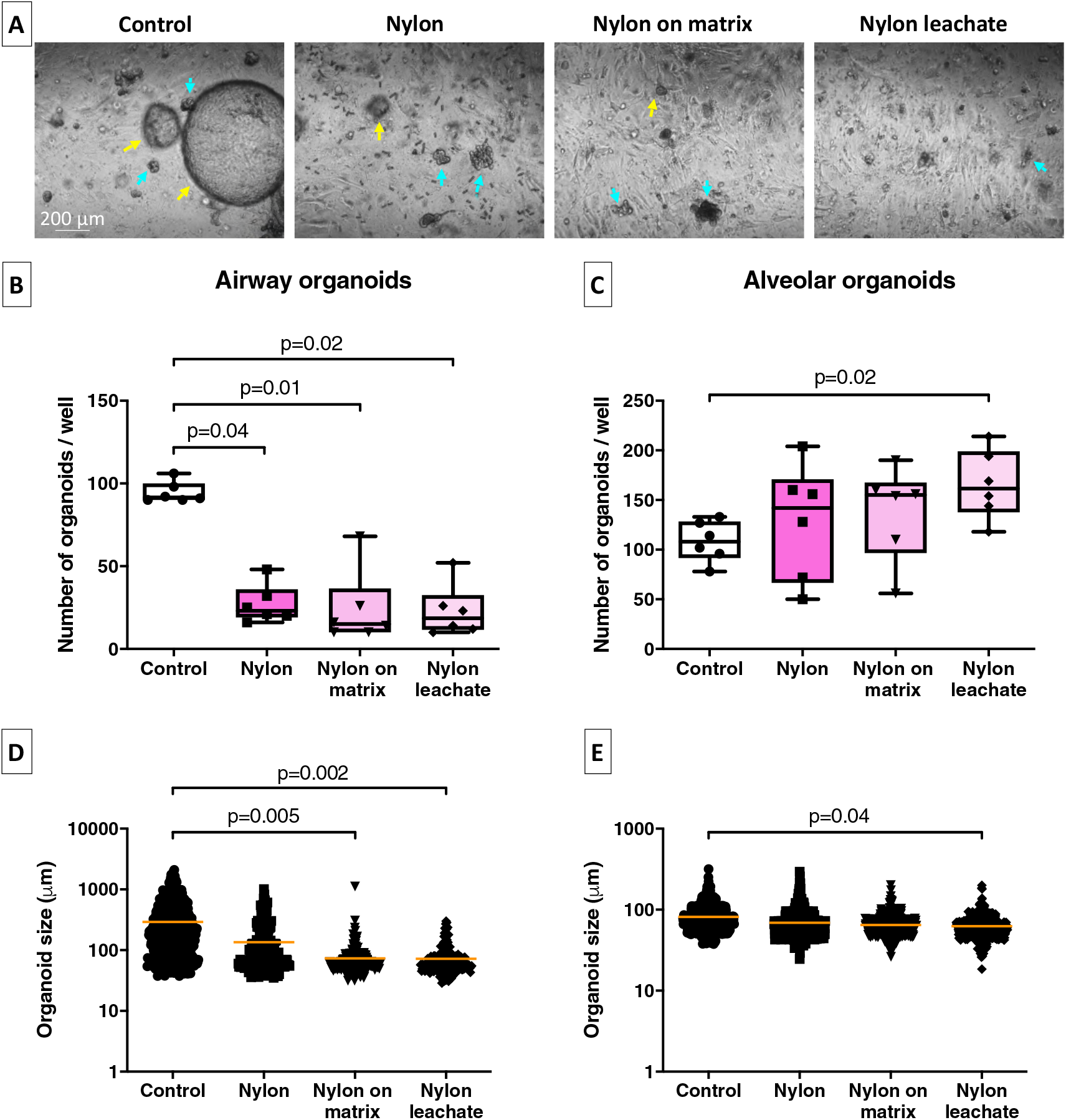
Impact of nylon reference microfibers and their leaching components on growth of murine lung organoids. (**A**) Representative light microscopy images of all treatment conditions. Yellow arrows in the light microscopy images indicate airway organoids, whereas cyan arrows indicate alveolar organoids. (**B and C**) Quantification of the numbers and (**D and E**) sizes of airway and alveolar organoids following either direct exposure to 5000 nylon microfibers or indirect exposure to nylon by adding 5000 microfibers (equivalent to 39 μg/ml nylon) on top of the Matrigel or by adding nylon leachate to the culture medium (n=6 independent isolations). Groups were compared using a Friedman test with Dunn’s correction for multiple testing. P<0.05 was considered significant.

The strong effects observed with the leachate suggested that some components and/or degradation products may leak and/or form during fiber ageing at 37C. Thus in order to determine the chemical identity of the components leaching from nylon reference microfibers we used mass spectrometry analysis. This revealed high concentrations of cyclic nylon oligomers (mono-, di- and trimers) in the leachate of nylon microfibers (Figure S4A), which are known to develop as by-products during the production of nylon (*44*). However, when exposing murine lung organoids to different concentrations of these isolated oligomers separately or in combination, we observed no effects on either number or size of organoids (Figure S4B-E showing the highest concentration that has been tested). These data suggest that other components in nylon leachate are causing the inhibitory effects on organoid growth. Recent work by Sait and Sørensen and colleagues showed that the most abundant chemicals leaching from nylon are bisphenol A and benzophenone-3 (*45, 46*). However, we could not detect these in our leachate, suggesting that if they are present, they are so in minute quantities. Initial experiments incubating organoids with different concentrations of bisphenol A or benzophenone-3 did not show effects on organoid growth of either of them suggesting they are indeed not the culprits in our leachate (data not shown).

### Leaching nylon components mainly affected developing organoids

As our experimental set-up specifically studied effects of microplastic fibers on developing organoids, we next studied whether already-developed, mature organoids were also affected by nylon microplastics. We therefore exposed organoids to nylon reference microfibers during organoid formation as before (14-day incubation) and we additionally exposed fully developed 14-day organoids to microfibers on top of the Matrigel or to nylon leachate for an additional 7 days. Interestingly, in contrast to the strong effects observed on developing organoids, we found that the compounds leaching from nylon had no effects on already-developed organoids (Figure 7A), as reflected by unchanged numbers of organoids (Figure 7B and C) and unchanged sizes (Figure 7D and E). This suggests that these nylon leachates are mostly harmful to differentiation of epithelial progenitors, but do not kill fully differentiated epithelial cells.

**Figure 7.**
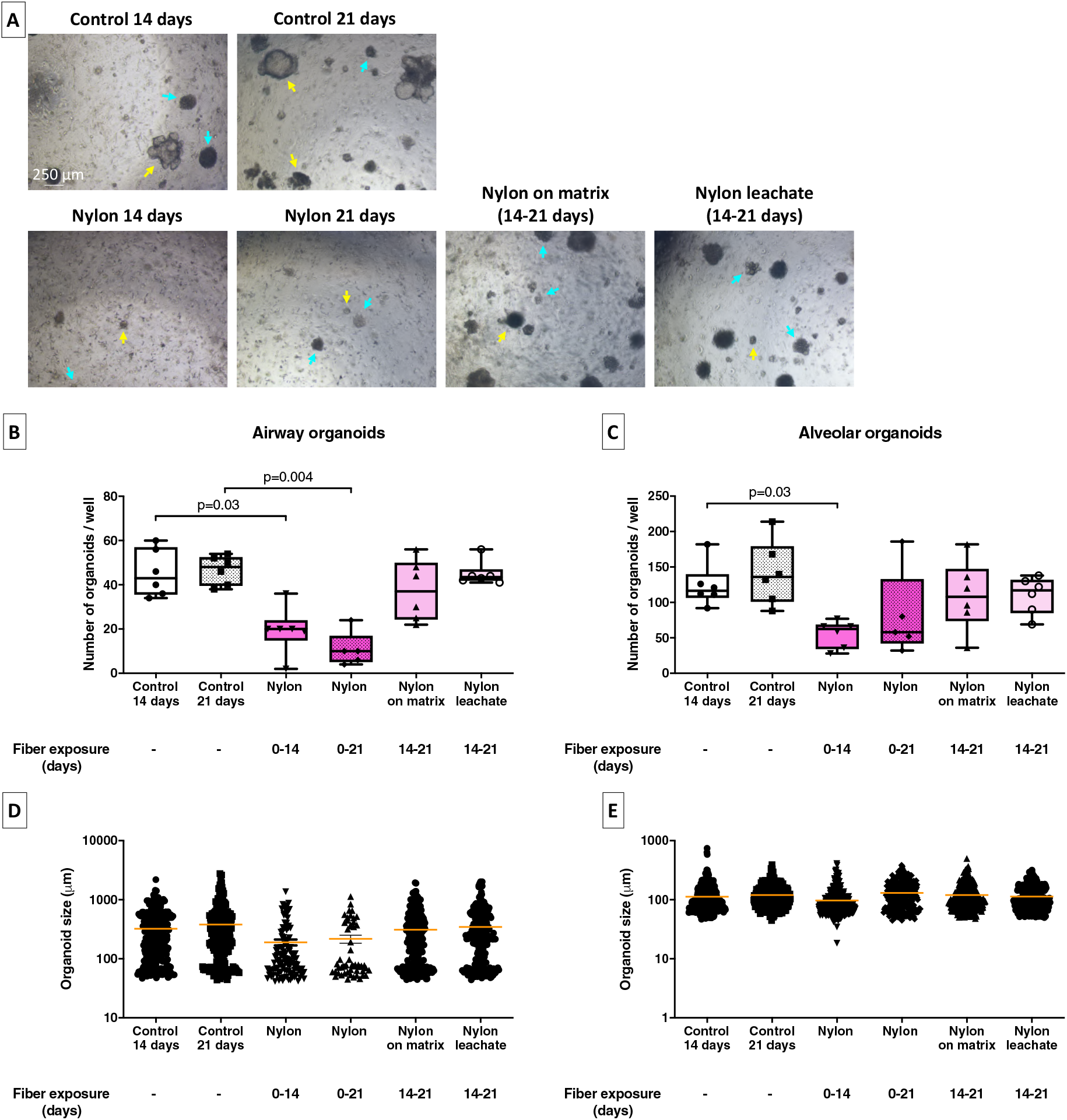
Effects of nylon reference fiber leachate on already-developed lung organoids. (**A**) Representative light microscopy images of all treatment conditions. Yellow arrows in the light microscopy images indicate airway organoids, whereas cyan arrows indicate alveolar organoids. (**B and C**) Quantification of the numbers and (**D and E**) sizes of airway and alveolar organoids following exposure to no or 5000 nylon microfibers (equivalent to 39 μg/ml nylon) for 14 or 21 days. A set of other organoids developed without treatment for 14 days and were exposed to nylon by adding 5000 microfibers (equivalent to 39 μg/ml nylon) on top of the Matrigel or by adding nylon leachate to the culture medium for another 7 days (n=6 independent isolations). Groups were compared using a Kruskal-Wallis test with Dunn’s correction for multiple testing. P<0.05 was considered significant.

### Exposure to nylon inhibited epithelial development pathways and stimulated expression of ribosome components

To better understand the mechanisms behind the observed effects on the growth of airway organoids, we performed bulk RNA-sequencing (RNAseq) analysis on epithelial cells and fibroblasts resorted from organoid cultures exposed to two different concentrations of nylon fibers (2000 or 5000 fibers) or not. The condition of 2000 fibers was added because the effect of 5000 fibers on airway epithelial development was already profound and we wanted to investigate more subtle changes. However, both conditions had an enormous impact on epithelial gene expression as depicted by the volcano plots (Figure 8A-D). Exposure to 2000 nylon fibers (equivalent to 16 μg/ml nylon) resulted in 16455 transcripts being differentially expressed at least two-fold compared to nonexposed controls, with an q value <0.05 (p value corrected for the false discovery rate), with most being downregulated (Figure 8A and C). Exposure to 5000 nylon fibers (equivalent to 39 μg/ml nylon) resulted in 39395 transcripts being differentially expressed at least two-fold compared to nonexposed controls, with most being upregulated (Figure 8B and D).

**Figure 8:**
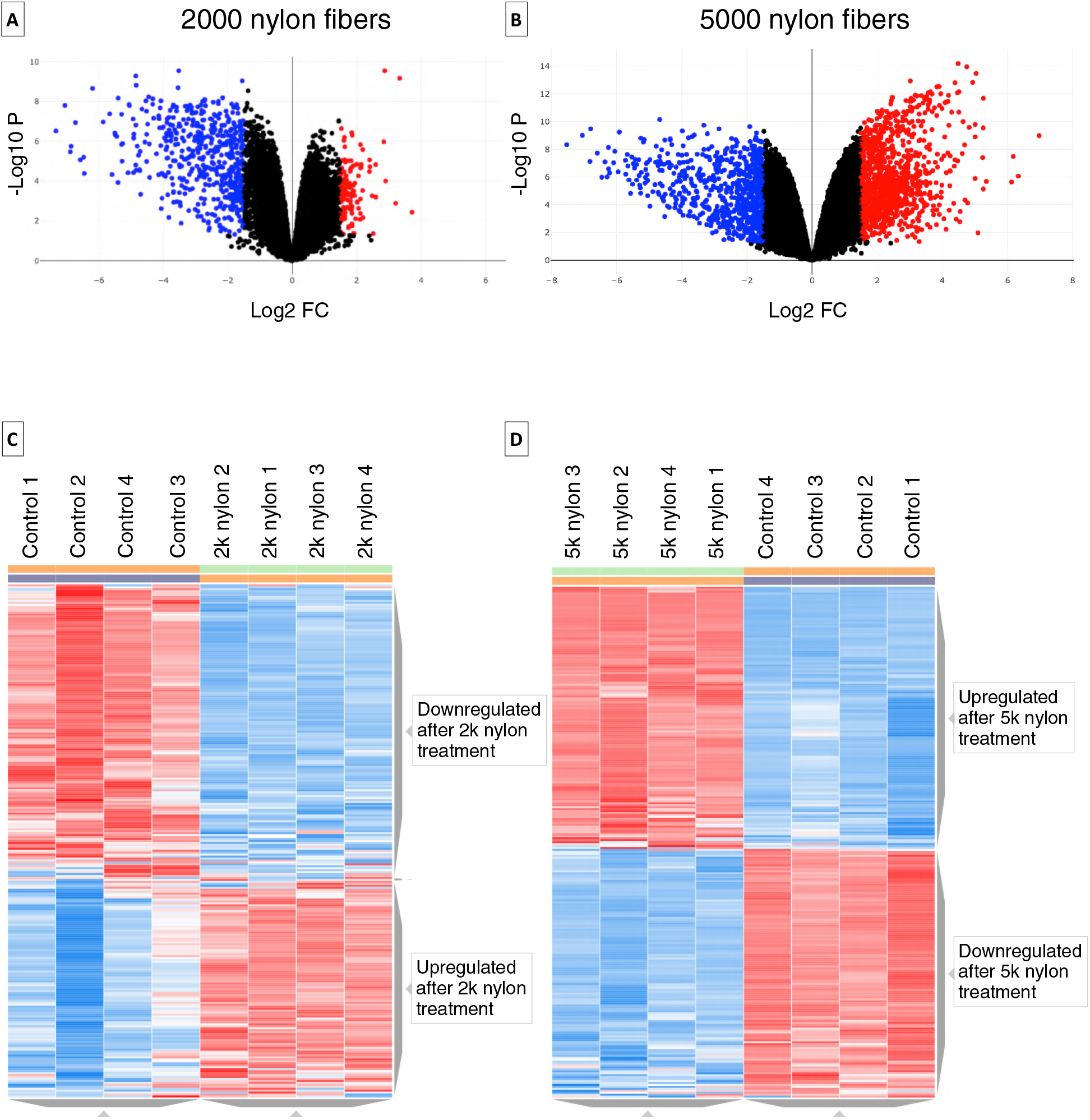
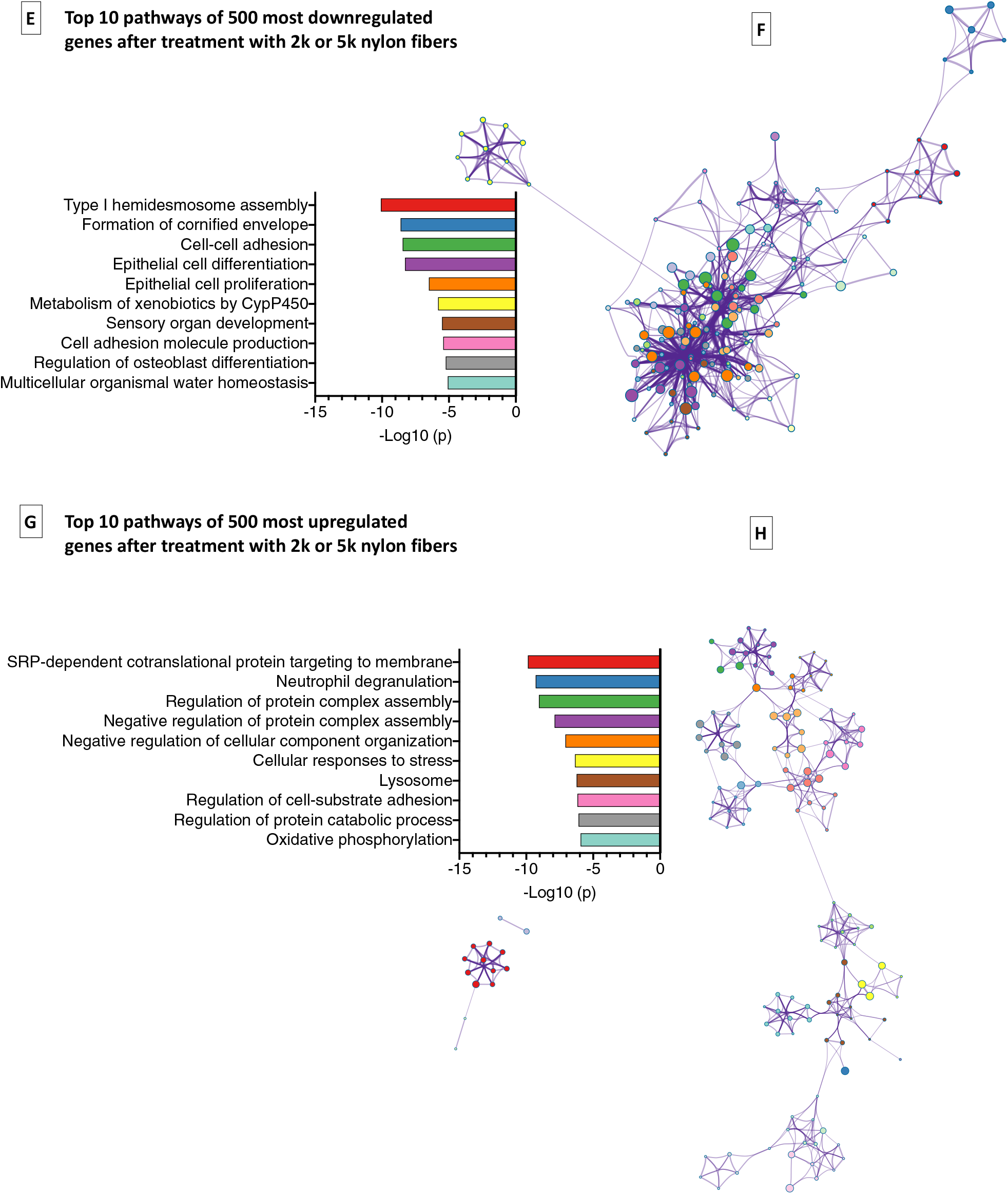
RNAseq analysis of epithelial cells exposed to nylon or not. (**A**) Volcano plot of differentially expressed genes by epithelial cells exposed to 2000 nylon fibers (equivalent to 16 μg/ml nylon) or not. (**B**) Volcano plot of differentially expressed genes by epithelial cells exposed to 5000 nylon fibers (equivalent to 39 μg/ml nylon) or not. Upregulated genes are marked in red, downregulated genes in blue. Genes were selected with thresholds of fold change >2 and q<0.05. (**C**) Unsupervised clustering heat map of epithelial cells exposed to 2000 nylon fibers (equivalent to 16 μg/ml nylon) or not. (**D**) Unsupervised clustering heat map of epithelial cells exposed to 5000 nylon fibers (equivalent to 39 μg/ml nylon) or not. (E) Metascape bar graphs of top 10 nonredundant enrichment clusters of genes downregulated by exposure to nylon ordered based on statistical significance (p value). (**F**) Metascape enrichment network visualization showing the intra-cluster and inter-cluster similarities of enriched terms of genes downregulated by exposure to nylon, up to ten terms per cluster. Cluster annotations colors are shown in bar graph of panel E. (**G**) Metascape bar graphs of top 10 nonredundant enrichment clusters of genes upregulated by exposure to nylon ordered based on statistical significance (p value). (**H**) Metascape enrichment network visualization showing the intra-cluster and inter-cluster similarities of enriched terms of genes upregulated by exposure to nylon, up to ten terms per cluster. Cluster annotations colors are shown in bar graph of panel E. 2k: 2000 fibers; 5k: 5000 fibers.

To reduce the number of transcripts, we then selected only those transcripts that had an average basemean expression of at least 10 and were significantly (q value <0.05) up or downregulated in both exposure conditions of 2000 and 5000 nylon fibers compared to nonexposed controls. This resulted in 10764 transcripts that were differentially expressed compared to nonexposed controls, with 5522 being downregulated and 5242 being upregulated. The downregulated transcripts were then sorted on the lowest q value for exposure to 2000 fibers and the upregulated transcripts were sorted on the lowest q value for exposure to 5000 fibers and the top 500 genes of each were used for pathway analysis using Metascape (*47*).

The top pathways identified for downregulated genes following nylon exposure were highly enriched for epithelial development and function (figure 8E-F), while the top pathways identified for upregulated genes were highly enriched for mRNA translation and protein synthesis (figure 8G-H).

We then investigated expression of individual genes in the top 5 enriched pathways for up and downregulated genes in more detail (Figures 8E-H). The top 5 enriched pathways for downregulated genes were type I hemidesmosome assembly, formation of cornified envelope, cell-cell adhesion, epithelial cell differentiation, and epithelial cell proliferation, while the top 5 pathways for upregulated genes were SRP-dependent cotranslational protein targeting to membrane, neutrophil degranulation, regulation of protein complex assembly, negative regulation of protein complex assembly, and negative regulation of cellular component organization.

We first investigated the downregulated genes and many of them represent important epithelial populations in the lung. We therefore investigated genes associated with specific epithelial populations (listed in table 1). The expression of these genes correlated well with our organoid findings that airway epithelial cell growth was most affected by exposure to nylon fibers, while alveolar epithelial cell growth was less affected (Figure 9). Both AECI and AEC II genes (Figure 9A) were only marginally lower expressed after exposure to nylon while most genes for basal cells (Figure 9B), ciliated cells (Figure 9C), club cells and goblet cells (Figure 9D) were expressed at significantly lower levels compared to controls with two noticeable exceptions: ciliated cell marker *Tuba1a* and club cell marker *Scgb1a1*, that were expressed at significantly higher levels compared to nonexposed controls. Proliferation markers like proliferation marker protein 67 (*Mki67*), forkhead box protein M1 (*Foxm1*), and polo-like kinase 1 (*Plk1*, Figure 9E) confirmed this general lack of proliferation in epithelial cells as all three were expressed at lower levels in a dose-dependent manner after exposure to nylon fibers. Expression of genes for signaling molecules essential for epithelial growth and development were impressively and dose-dependently downregulated by nylon exposure as well (Figure 9F), including *Notch1* and *Notch2* and their ligands Jagged 1 (*Jag1*) and 2 (*Jag2*) (*48–50*), *Bmp4* and *Bmp7* (*51–53*), *Wnt4* and *Wnt7a* (*54, 55*), and the receptor for hepatocyte growth factor, *Met*. The lower expression of basal cell-specific markers and essential factors that are needed for differentiation of other cell types like ciliated and goblet cells may explain why the growth of in particular airway organoids was inhibited most by nylon.

**Figure 9:**
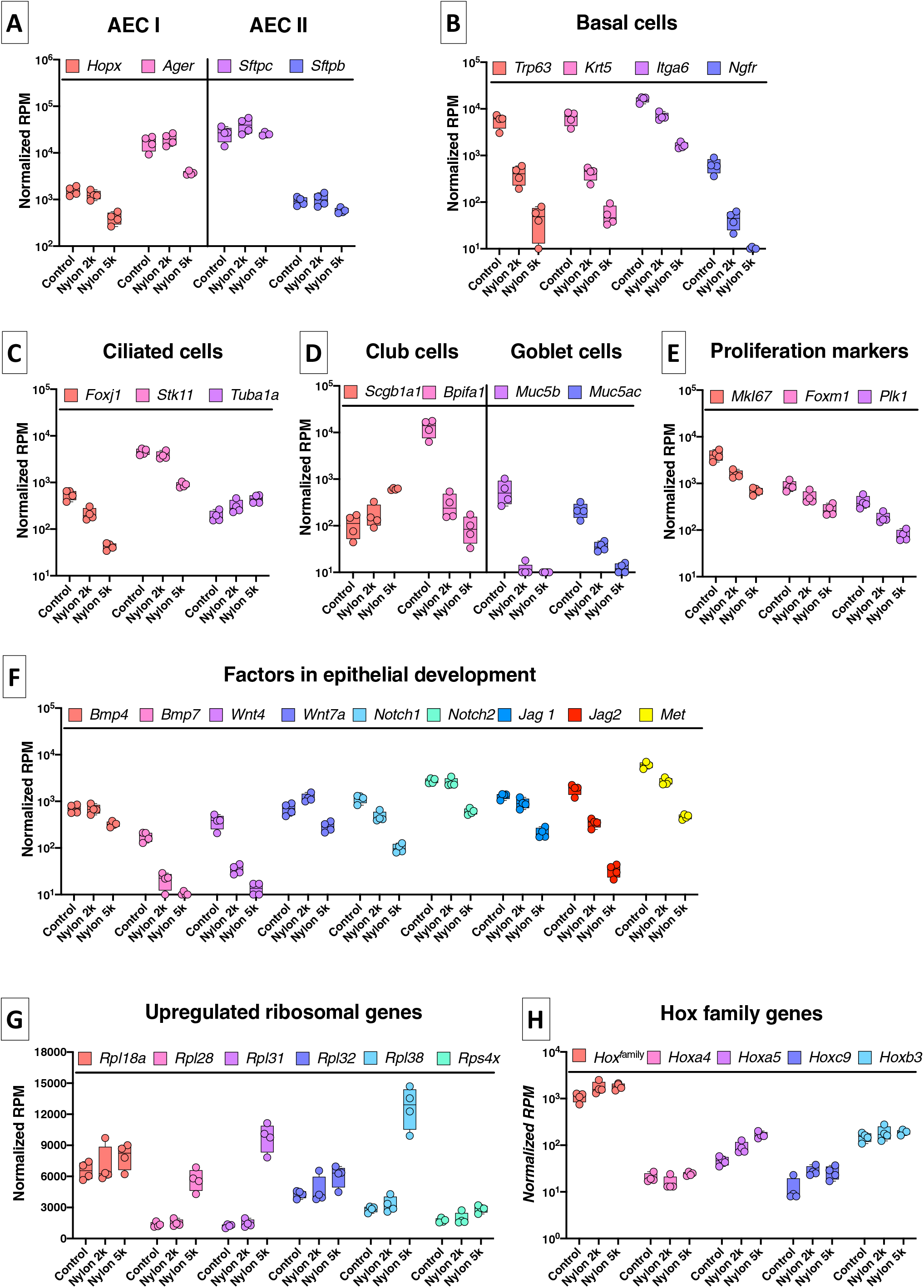
Expression profiles of individual genes from the pathway analyses. Genes shown were significantly differentially expressed in epithelial cells isolated from organoids exposed to 2000 (2k) or 5000 (5k) nylon fibers compared to untreated controls according to a false discovery rate of q<0.05 (n=4 independent isolations). The following genes in the following conditions were not significantly different: 2k nylon: Hopx, Ager, Sftpc, Sftpb, Stk11, Scgb1a1, Bmp4, Notch2, Hoxa4; 5k nylon: Hopx, Ager, Plk1. (A) Genes highly expressed by alveolar epithelial cells type I (AECI) and type II (AECII). (**B**) Genes highly expressed in basal cells. (**C**) Genes highly expressed in ciliated cells. (**D**) Genes highly expressed in club cells and goblet cells. (**E**) Genes associated with proliferation. (**F**) Genes encoding factors important for epithelial development. (**G**) Genes encoding ribosomal proteins. (**H**) Genes encoding Hox family genes.

**Table 1:** Markers associated with different epithelial populations in lung tissue.

| Cell type | Protein name | Gene name |
| --- | --- | --- |
| **Epithelial cells present in airways** |  |  |
| Basal cells | Transformation-related protein 63 | *Trp63* |
|  | Keratin 5 | *Krt5* |
|  | Integrin alpha-6 | *Itga6* |
|  | Nerve growth factor receptor | *Ngfr* |
| Ciliated epithelial cells | Forkhead box J1 | *Foxj1* |
|  | Serine/threonine kinase 11 | *Stk11* |
|  | Acetylated alpha tubulin | *Tuba1a* |
| Goblet cells | Mucin 5 subtype B | *Muc5b* |
|  | Mucin subtype AC | *Muc5ac* |
| Club cells | Secretoglobin family 1A member 1 | *Scgb1a1* |
|  | BPI fold containing family A member 1 | *Bpifa1* |
| **Epithelial cells present in alveoli** |  |  |
| Alveolar epithelial cells type I (AECI) | Homeodomain-only protein homeobox | *Hopx* |
|  | Advanced glycosylation end-product specific receptor | *Ager* |
| Alveolar epithelial cells type II (AECII) | Surfactant protein C | *Sftpc* |
|  | Surfactant protein B | *Sftpb* |

To exclude the possibility that these effects on epithelial cells were the result of a decreased support function from fibroblasts, for instance by nylon selectively killing fibroblasts or inhibiting the expression of important growth factors, we separately analyzed the resorted fibroblast fraction for expression of proliferation genes and important growth factors, i.e. *Mki67*, *Foxm1*, *Plk1*, fibroblast growth factors 2, 7, and 10 (*Fgf2*, *Fgf7*, and *Fgf10*), *Wnt2*, *Wnt5a*, and hepatocyte growth factor (*Hgf*) (*33*). None of these genes were negatively affected in fibroblasts after exposure to nylon fibers compared to untreated controls and expression of most actually went up slightly (Figure S5).

The genes most prominently upregulated after exposure to nylon fibers were mostly encoding for ribosomal proteins, with ribosomal protein L28 (*Rpl28*), L31 (*Rpl31*), and L38 (*Rpl38*) being most profoundly upregulated (Figure 9G). Ribosomal proteins are a large family of proteins and essential parts of ribosomes translating mRNA to protein. Recent data has shown that heterogeneity in ribosomal protein composition within ribosomes makes them selective for translating subpools of transcripts (*56*). For instance, Rpl38 has been shown to regulate translation of the homeobox (*Hox*) genes, that are key in anatomical development (*57*). We therefore also investigated the expression of all *Hox* genes and some specific members of this family involved in lung epithelial development in more detail (Figure 9H) (*58, 59*). After exposure to nylon fibers, expression of Hox family members was significantly higher and this pattern was seen for highlighted members *Hoxa4*, *Hoxa5*, *Hoxc9*, and *Hoxb3* as well.

## Discussion

Recent reports have shown that man-made fibers are ubiquitously present in indoor air (*9, 35–38*). It is estimated that approximately 30% of those indoor fibers are of plastic origin, particularly from textiles (*9, 35–38*). The lungs are continuously exposed to this airborne microplastic pollution (*60*), but the consequences of common household exposure on our lungs are unclear. In our present work, we found that both polyester and nylon microfibers negatively affected the growth and development of human and murine lung organoids, with nylon being the most harmful. Already established lung organoids were not affected and therefore our results may be of particular importance for young children with developing airways and for people undergoing high levels of epithelial repair, such as people with respiratory diseases.

Nylon was found to be the most consistently harmful for growth of airway organoids and was less inhibitory for growth of alveolar organoids, while polyester affected both types equally but less profoundly than nylon. This was the case for our reference fibers as well as environmentally relevant fibers made from fabrics purchased in a local fabric store. The gene expression analysis also confirmed that growth of alveolar organoids was affected less by nylon exposure and even appeared *induced* after treatment with leachate or lower numbers of nylon fibers. The explanation for this finding may be found in the downregulation of Notch signaling pathway members by nylon fibers. Multiple studies have shown that Notch signaling is required for development of airway epithelial cells, most specifically goblet cells, whereas disruption of this signaling boosts alveolar epithelial development (*61, 62*). Both *Notch1* and *Notch2*, as well as their ligands *Jag1* and *Jag2* were expressed at significantly lower levels after nylon treatment, suggesting disruption of Notch signaling may be responsible for the divergent effect nylon has on airway versus alveolar epithelial growth.

The Notch pathway, incidentally, is also important for the development of club cells (*63*). Morimoto and colleagues showed that *Notch2* was involved in the decision between club cell or ciliated cell development. Interestingly, the gene expression data for club cells and ciliated cells were somewhat ambiguous, with important markers for these cell types having higher expression (*Scgb1a1* for club cells and *Tub1a1* for ciliated cells) and others having lower expression (*Bpifa1* for club cells and *Foxj1* and *Stk11* for ciliated cells) after nylon exposure. This may suggest that the lower expression of Notch2 may be hampering the cell fate decision between club and ciliated cells in some way, resulting in improperly differentiated cell types. In combination with the inhibited development of basal epithelial cells and goblet cells, this altered airway epithelial differentiation may explain the bronchiolitis found in nylon flock workers and rats exposed to nylon (*64–66*).

Our results demonstrated that the negative effect of nylon fibers on development of particularly airway organoids was caused by components leaching from these fibers. As the most abundant components in this leachate, the cyclic oligomers, were not responsible for this effect, we used RNAseq analysis to look for signatures of other components. Although we could not detect bisphenol A in our leachate, it is of interest to note that exposure of fruit flies to bisphenol A specifically upregulated ribosome-associated genes (*67*), similar to what we found in our study. The association of these *Rpl* genes with the *Hox* family genes is of particular interest for lung epithelial development. *Rpl38* was one of the three most upregulated *Rpl* genes by nylon and was shown to interact with a specific subset of *Hox* genes including *Hoxa4*, *Hoxa5*, and *Hoxb3* (*57*). These have all been associated with epithelial differentiation, with *Hoxa5* taking a center stage in goblet versus club cell differentiation (*58, 59*). Boucherat and colleagues showed that loss of *Hoxa5* drove epithelial differentiation towards goblet cell differentiation at the expense of club cell differentiation and *Scgb1a1* expression (*58*). Remarkably, *Hoxa5* was expressed at significantly higher levels in our dataset in a dose-dependent manner after exposure to nylon and we also found a matching dose-dependent increase in *Scgb1a1* and decrease in *Muc5ac* and *Muc5b* expression. Taken together our data suggest that component (s) in nylon leachate may be skewing differentiation of epithelial progenitors away from airway epithelial cells possibly through changes in expression of *Hoxa5* and/or *Notch*. Which components or combinations of components are responsible for these effects is still an open question.

A strength of using lung organoids is the opportunity to directly translate murine findings to human lung epithelial repair (*32, 33, 43, 68, 69*). Using cells isolated from human lungs we have shown human epithelial cells respond similarly to polyester and nylon fibers, demonstrating our results are relevant for human epithelial differentiation and growth too. Despite this advantage, lung organoids are a relatively simple model of lung tissue and lack the immune and endothelial compartment present *in vivo*. Especially having the immune compartment present could alter how lung tissue responds to these microplastic fibers. For example, innate immune cells like macrophages are also one of the first cells to come into contact with microplastic fibers following inhalation and macrophages are known to respond strongly to inhaled particles and fibers (*70*) and are also important for lung repair (*71*). It is therefore recommended to include lung macrophages in lung organoid cultures in future studies as was done before by Choi *et al.* (*72*). This way, a more comprehensive view on the interaction between pivotal lung cells and microplastics can be obtained.

The implication of our results for the human population is of high relevance. It is important to note that similar to the high occupational exposure in industry workers, the microplastic fiber doses as used in our *in vitro* experiments are much higher than daily exposure for most people. Previous studies estimated that a male person with light activity may inhale around 272 microplastic particles per day based on air sampling using a breathing thermal manikin (*60*). The total surface of airway epithelial cells is 2471 cm^2^ for human lungs (*73*), resulting in 0.1 particle/cm^2^. Our murine cultures had an average of 70 airway organoids with an average size of 320 μm, resulting in a total surface of 0.2 cm^2^ that was incubated with 5000 fibers or 25,000 fibers/cm^2^. For our calculations we assumed these particles/fibers will get trapped onto airway epithelial cells. Generally particles or fibers of sizes between 10 and 100 µm will deposit onto epithelial cells covering airway walls (*40*). Only fibers with a diameter smaller than 3 µm have the ability to penetrate deep into the lungs and reach the alveoli. Our fiber sizes were limited by the availability of polyester and nylon filaments of standardized small diameters and therefore were not small enough for alveolar deposition. However, airway trapping can still cause local harm as we found nylon fibers to inhibit airway epithelial differentiation most. It will therefore be crucial to study in more detail how many and what kind of fibers deposit in which regions of the lungs and what fraction can still be cleared. In addition, we need to gather more information about exposure levels in indoor environments to assess real-life inhalation levels. A limitation here is the detection of microplastics, as smaller particles might escape current detection methods (*10, 74*). Especially our finding that epithelial differentiation and repair mechanisms are affected most by microplastics exposure, suggests airborne microplastics may be most harmful to young children with developing airways and to people undergoing high levels of epithelial repair. These could be people with a chronic lung disease or even healthy individuals suffering from a seasonal respiratory virus infection.

In conclusion, with the ongoing and growing use of plastics, potential associated health problems in the human population may also increase. The results of the present study strongly encourage to look in more detail at both hazard of and exposure to microplastic fibers, and outcomes of these experiments will be valuable to advise organizations such as the World Health Organization and Science Advice for Policy by European Academies who have recently stated that more research is urgently needed (*30, 31*). Importantly, future research should focus on examining the presence and number of such fibers both in our indoor environment and in human lung tissue, to better estimate the actual risk of these fibers to human health.

## Materials and Methods

### Production of microfibers and leachate

#### Reference microfibers and leachate

Microfibers of standardized dimensions were produced as described before (*34*). In short, polyester and nylon fibers (both Goodfellow, UK) with filament diameters of 14±3.5 μm and 10±2.5 μm respectively were aligned by wrapping them around a custom-made spool, coated with a thin layer of cryo compound (KP-CryoCompound, VWR International B.V., PA, USA) and frozen. Aligned fibers were cut into similar length parts (∼2 cm) using a scalpel (Swann-Morton, UK) and moulded onto a compact block that was oriented perpendicular to the base of a cryomicrotome (Microm HM 525, Thermo Fisher Scientific, MA, USA). Microfibers were cut at lengths of 50 μm for polyester and 30 μm for nylon, after which the fibers were thawed, washed with water through a 120 μm filter (Merck Millipore, MA, USA) to remove miscut fibers and contaminants, collected by vacuum filtration using 8 μm polycarbonate membrane filters (Sterlitech, WA, USA) and stored dry at −20°C.

Nylon leachate was produced by incubation of nylon reference microfibers in phosphate buffered saline (PBS) for 7 days at 37°C in the dark, followed by filtration using a 0.2 μm syringe filter (GE Healthcare Life Sciences, UK). The leachate was stored at −20°C until further use.

#### Environmental microfibers

Environmental polyester and nylon textile microfibers were prepared from commercially available pure fabrics. White polyester fabric was washed at 40°C in a washing machine (Samsung, South Korea) and dried in a tumble dryer (Whirlpool, MI, US). Fibers with an estimated filament diameter of 15 μm were collected on the filter of the tumble dryer and subsequently frozen with cryo compound and sectioned into lengths of 50 μm using a cryomicrotome. White nylon fabric (estimated filament diameter of 40 μm) was cut into small squares, stacked, frozen with cryo compound, and cut into lengths of 12 μm. All microfibers were thawed, washed with water through a 120 μm filter, collected by vacuum filtration (8 μm filter) and finally stored at −20°C.

### Characterization of microfibers and leachate

#### Scanning electron microscopy

Samples were prepared for scanning electron microscopy (SEM) analysis on an aluminium sample holder using adhesive carbon-coated tape. Excessive microfibers were removed using pressurized air, after which the samples were sputter-coated with 10 nm of gold. Images were obtained using a JSM-6460 microscope (Jeol, Japan) at an acceleration voltage of 10 kV.

#### Dimensions

Digital photomicrographs were captured at 200× magnification using a Nikon Eclipse TS100 inverted microscope coupled to a Nikon Digital Sight DS-U3 microscope camera controller (both Japan), after which microfiber diameters and lengths (median of 200) were determined using NIS-Elements AR 4.00.03 software.

### Ethics

#### Animal experiments

The experimental protocol for the use of mice for epithelial cell isolations was approved by the Animal Ethical Committee of the University of Groningen (The Netherlands) and all experiments were performed according to strict governmental and international guidelines on animal experimentation. C57BL/6 mice (8-14 weeks) were bred at the Central Animal Facility of the University Medical Center Groningen (UMCG) (IVD 15303-01-004). Animals received ad libitum normal diet with a 12 h light/dark cycle.

#### Human lung tissue

Histologically normal lung tissue was anonymously donated by individuals with COPD (n=6) or without COPD (n=1) undergoing surgery for lung cancer and not objecting to the use of their tissue. COPD patients included ex- and current-smoking individuals with GOLD stage I-IV disease (GOLD I=1, GOLD II=2, GOLD IV=3). Subjects with other lung diseases such as asthma, cystic fibrosis, or interstitial lung diseases were excluded. The study protocol was consistent with the Research Code of the University Medical Center Groningen (UMCG) and Dutch national ethical and professional guidelines (www.federa.org). Sections of lung tissue of each patient were stained with a standard haematoxylin and eosin staining and checked for abnormalities by a lung pathologist.

### Cell cultures

Mouse lung fibroblasts (CCL-206, ATCC, Wesel, Germany) or human lung fibroblasts (MRC5, ATCC, CCL-171) were cultured in 1:1 DMEM (Gibco, MD, USA) and Ham’s F12 (Lonza, Switzerland) or Ham’s F12 respectively, both supplemented with 10% heat inactivated fetal bovine serum (FBS, GE Healthcare Life Sciences), 100 U/ml penicillin and 100 μg/ml streptomycin, 2 mM L-glutamine and 2.5 μg/ml amphotericin B (all Gibco).

Fibroblasts were cultured at 37°C under 5% CO_2_ and humidified conditions. For use in organoid cultures, near-confluent cells were proliferation-inactivated by incubation with 10 μg/ml mitomycin C (Sigma-Aldrich, MO, USA) in cell culture medium for 2 hours, after which they were washed in PBS, and left in normal medium for at least 1 hour before trypsinizing and counting.

### Lung epithelial cell isolation

#### Mouse lung epithelial cell isolation

Epithelial cells were isolated from lungs of mice using antibody-coupled magnetic beads (microbeads) as described before (*32, 33, 68*). In short, mice were sacrificed under anaesthesia, after which the lungs were flushed with 0.9% NaCl and instilled with 75 caseinolytic units/1.5 ml dispase (Corning, NY, USA). After 45 minutes of incubation at room temperature, the lobes (excluding trachea and extrapulmonary airways) were homogenized in DMEM containing 100 U/ml penicillin and 100 μg/ml streptomycin and 40 μg/ml DNase1 (PanReac AppliChem, Germany), washed in DMEM (containing penicillin, streptomycin and DNase1), and the digested tissue was passed through a 100 µm cell strainer. The suspension was incubated for 20 minutes at 4°C with anti-CD31 and anti-CD45 microbeads in MACS buffer and magnetically sorted using LS columns. The CD31−/CD45− flow-through was incubated for 20 minutes at 4°C with anti-CD326 (EpCAM) microbeads in MACS buffer, after which purified epithelial lung cells were obtained by positive selection using LS columns. CD326-positive epithelial cells were resuspended in CCL206 fibroblast medium, counted and seeded into Matrigel immediately after isolation with equal numbers of CCL206 fibroblasts, as described below. All materials were purchased at Miltenyi Biotec (Germany) unless stated otherwise.

#### Human lung epithelial cell isolation

Human lung epithelial cells were isolated from lung tissue specimens obtained from patients. Peripheral lung tissue was minced and dissociated in DMEM-containing enzymes (Multi Tissue Dissociation Kit) at 37°C using a gentleMACS Octo Dissociator. The cell suspension was filtered (70 µm and 35 µm nylon strainer, respectively) prior to 20-minute incubation at 4°C with anti-CD31 and anti-CD45 microbeads in MACS buffer. The CD31−/CD45− fraction was obtained by negative selection using an AutoMACS. Epithelial cells were then isolated by positive selection after 20-minute incubation at 4°C with anti-CD326 (EpCAM) microbeads in MACS buffer. Human EpCAM+ cells were resuspended in 1:1 DMEM and Ham’s F12, supplemented with 10% heat inactivated FBS, 100 U/ml penicillin and 100 μg/ml streptomycin, 2 mM L-glutamine and 2.5 μg/ml amphotericin B, counted and seeded into Matrigel immediately after isolation with equal numbers of MRC5 fibroblasts. All materials were purchased at Miltenyi Biotec unless stated otherwise.

### Lung organoid cultures

Lung organoids were grown as previously described with minor modifications (*32, 33, 68*). For mouse lung organoids, 10,000 EpCAM+ cells and 10,000 CCL206 fibroblasts were seeded, and for human lung organoids, 5,000 EpCAM+ cells and 5,000 MRC5 fibroblasts were seeded in 100 μl growth factor-reduced Matrigel matrix (Corning) diluted 1:1 in DMEM:Ham’s F-12 1:1 containing 10% FBS, 100 U/ml penicillin and 100 μg/ml streptomycin, 2 mM L-glutamine and 2.5 μg/ml amphotericin B into transwell 24-well cell culture plate inserts (Corning). Organoids were cultured in DMEM:Ham’s F-12 1:1 supplemented with 5% FBS, 100 U/ml penicillin and 100 μg/ml streptomycin, 2 mM L-glutamine, 2.5 μg/ml amphotericin B, 4 ml/l insulin-transferrin-selenium (Gibco), 25 μg/l recombinant mouse (Sigma-Aldrich) or human epithelial growth factor (EGF, Gibco) and 300 μg/l bovine pituitary extract (Sigma-Aldrich). During the first 48h of culture, 10 μM Y-27632 (Axon Medchem, the Netherlands) was added to the medium.

A titration curve for polyester or nylon microfibers was made with 2000, 3000, 4000 or 5000 fibers per well corresponding to approximately 49, 73, 98 and 122 μg/ml of polyester fibers or 16, 23, 31, and 39 μg/ml of nylon fibers. For all other experiments, 5000 polyester or nylon reference (equivalent to 122 μg/ml of polyester or 39 μg/ml of nylon) or environmental microfibers (equivalent to 122 μg/ml of polyester or 39 μg/ml of nylon and) were used per condition. Fibers were in direct contact with developing organoids during 14 days by mixing them with Matrigel and cells prior to seeding in the insert, except for those experiments studying effects of leaching nylon components. In those cases, 5000 polyester and nylon reference fibers were added on top of the organoids, thereby excluding physical contact between the microfibers and the developing organoids, or equivalent amounts of fiber leachate were added to the medium during 14 days of organoid culture. For testing the effects of nylon oligomers, concentrations between 26.8 ng/ml and 53.6 µg/ml were used; the latter concentration being twice as high as the used fiber concentrations (5000 fibers per condition). Oligomers were synthesized and characterized as described in the supplementary materials and methods. All organoid cultures were maintained for 14 to 21 days at 37 °C under 5% CO_2_ and humidified conditions. Medium was refreshed 3 times per week.

Organoid colony forming efficiency was quantified by manually counting the total number of organoids per well after 14 or 21 days of culturing using a light microscope at 100× magnification. For mouse organoids, a distinction was made between airway and alveolar organoids, whereas for human organoids only one organoid phenotype was distinguished. The diameter of the organoids was measured using a light microscope (Nikon, Eclipse Ti), only including organoids larger than 50 µm in diameter, with a maximum of 50 organoids per phenotype per well.

### Immunofluorescence staining

Organoid cultures in Matrigel were washed with PBS, fixed in ice-cold 1:1 acetone and methanol (both Biosolve Chimie, France) for 15 minutes at −20°C and washed again with PBS after which aspecific antibody-binding was blocked for 2 hours in 5% bovine serum albumin (BSA, Sigma-Aldrich). Organoids were incubated overnight at 4°C with the primary antibodies (mouse anti-acetylated α-tubulin (Santa Cruz, TX, USA) and rabbit anti-prosurfactant protein C (Merck, Germany)) both diluted 1:200 in 0.1% BSA and 0.1% Triton (Sigma-Aldrich) in PBS. Next, after extensive but gentle washing with PBS, organoids were incubated with the appropriate Alexa-conjugated secondary antibody for 2 hours at room temperature (Alexa Fluor 488 donkey anti rabbit IgG and Alexa Fluor 568 donkey anti mouse IgG, both Thermo Fisher Scientific) diluted 1:200 in 0.1% BSA and 0.1% Triton in PBS. Organoid cultures were washed with PBS, excised using a scalpel, mounted on glass slides (Knittel, Germany) using mounting medium containing DAPI (Sigma-Aldrich) and covered with a cover glass (VWR). Digital photomicrographs were captured at 200× magnification using a DM4000b fluorescence microscope and LAS V4.3 software (both Leica, Germany).

### Isolation of epithelial cells and fibroblasts from organoid cultures

200,000 mouse EpCAM+ primary cells and 200,000 CCL206 fibroblasts (n=4 independent isolations) were seeded in 1 ml Matrigel diluted 1:1 in DMEM containing 10% FBS in 6-well plates (Greiner Bio-One, The Netherlands). 12.000 or 30.000 nylon reference microfibers were mixed with Matrigel and cells prior to seeding. Murine organoid culture medium was maintained on top and refreshed every two days. After 7 days, organoid cultures were digested with 50 caseinolytic units/ml dispase for 30 minutes at 37°C, transferred to 15 ml tubes, washed with MACS BSA stock solution and autoMACS rinsing solution (both Miltenyi), and digested further with trypsin (VWR) diluted 1:5 in PBS for 5 minutes at 37°C. The cell suspension was then incubated for 20 minutes at 4°C with anti-EpCAM microbeads in MACS buffer, after which the suspension was passed through LS columns. Both the EpCAM+ (epithelial cells) and EpCAM− (fibroblasts) cell fractions were used for RNA isolation and subsequent sequencing.

### Library preparation and RNA sequencing

Total RNA was isolated from EpCAM+ and EPCAM- cell fractions using a Maxwell® LEV simply RNA Cells/Tissue kit (Promega, WI, USA) according to manufacturer’s instructions. RNA concentrations were determined using a NanoDrop One spectrophotometer (Thermo Fisher Scientific). Total RNA (300 ng) was used for library preparation. Paired-end sequencing was performed using a NextSeq 500 machine (Illumina; mate 1 up to 74 cycles and mate 2 up to 9 cycles). Mate 1 contained the first STL (stochastic labeling) barcode, followed by the first bases of the sequenced fragments, and mate 2 only contained the second STL barcode. The generated data were subsequently demultiplexed using sample-specific barcodes and changed into fastq files using bcl2fastq (Illumina; version 1.8.4). The quality of the data was assessed using FastQC (*75*). The STL barcodes of the first mate were separated from the sequenced fragments using an in-house Perl script. Low quality bases and parts of adapter sequences were removed with Cutadapt (version 1.12; settings: q=15, O=5, e=0.1, m=36) (*76*). Sequenced poly A tails were removed as well, by using a poly T sequence as adapter sequence (T (100); reverse complement after sequencing). Reads shorter than 36 bases were discarded. The trimmed fragment sequences were subsequently aligned to all known murine cDNA sequences using HISAT2 (version 2.1.0; settings: k=1000, --norc). The number of reported alignments, k, was given a high number in order to not miss any alignment results (some genes have up to 62 transcripts). Reads were only mapped to the forward strand (directional sequencing). Fragment sequences that mapped to multiple genes were removed (unknown origin). When fragments mapped to multiple transcripts from the same gene all but one were given a non-primary alignment flag by HISAT2 (flag 256). These flags were removed (subtraction of 256) by the same Perl script in order to be able to use the Bash-based shell script (dqRNASeq; see below) that is provided by Bioo Scientific (Perkin Elmer, MA, USA). Fragments that mapped to multiple transcripts from the same gene were considered unique and were counted for each of the transcripts. The number of unique fragments (or read pairs) was determined for each transcript using the script provided by Bioo Scientific (dqRNASeq; settings: s=8, q=0, m=1). Counts that were used for further analysis were based on a unique combination of start and stop positions and barcodes (USS + STL). The full data set is available as Supplemental Table S3 (Supplemental Material available online; see https://figshare.com/s/26a93797d19154dc418a).

### Data analyses and statistics

All statistics were performed with GraphPad Prism 9·0. Nonparametric testing was used to compare groups in all experiments. For comparison of multiple-groups, a Kruskall wallis or Friedman test was used for nonpaired or paired nonparametric data respectively with Dunn’s correction for multiple testing. Differences in organoid size between groups were tested by using the average size of the organoids per independent experiment. Data are presented as median ± range and p-values <0·05 were considered significant.

For RNA sequencing data, the principal component analyses were performed in R using the R package DESeq2 (version 1.26.0) (*77*) in order to visualize the overall effect of experimental covariates as well as batch effects (function: plotPCA). Differential gene expression analyses (treated vs. nontreated) was performed with the same R package (default settings; Negative Binomial GLM fitting and Wald statistics; design=∼condition), following standard normalization procedures. Transcripts with differential expression >2 (nylon-treated versus nontreated fibroblasts or epithelial cells) and a false discovery rate smaller than 0.05 (q value) were considered differentially expressed in that specific cell type. Volcano plots and clustering heat maps were made using BioJupies (*78*). Pathway analysis was done using Metascape (*47*).

## Acknowledgments

### General

The authors thank Habibie (University of Groningen, Department of Molecular Pharmacology) for his help with the human organoid experiments, Imco Sibum, Paul Hagedoorn, and Anko Eissens (University of Groningen, Department of Pharmaceutical Technology and Biopharmacy) for their assistance at the scanning electron microscope, Andreas W. Ehlers (University of Amsterdam, Van ’t Hoff Institute for Molecular Sciences) for his assistance with the NMR spectroscopy, and Elena Höppener (TNO, Department Environmental Modeling Sensing and Analysis) for the energy dispersive X-ray and infrared spectroscopy analysis of the microfibers.

### Funding

ZonMW is gratefully acknowledged for their financial support with Microplastics and Health grant 458001013.

### Author contributions

FD: Study design, collection and assembly of data, data analysis and interpretation, manuscript writing, critical reading and revision.

SS: Collection and assembly of data, data analysis and interpretation, critical reading and revision.

GE: Collection and assembly of data, data analysis and interpretation, critical reading and revision.

XW: Collection and assembly of data, data analysis and interpretation, critical reading and revision.

IB: Collection and assembly of data, critical reading and revision.

DB: Collection and assembly of data, data analysis and interpretation, experimental material support, critical reading and revision.

IK: Collection and assembly of data, data analysis and interpretation, experimental material support, critical reading and revision.

DS: Collection and assembly of data, data analysis and interpretation, experimental material support, critical reading and revision.

RW: Collection and assembly of data, data analysis and interpretation, experimental material support, critical reading and revision.

MC: Collection and assembly of data, data analysis and interpretation, experimental material support, critical reading and revision.

AS: Data analysis and interpretation, critical reading and revision.

RG: Data analysis and interpretation, study design, experimental material support, critical reading and revision.

BM: Collection and assembly of data, study design, data analysis and interpretation, financial support, manuscript writing, critical reading and revision.

### Competing interests

The authors declare no competing interests.

## Data and materials availability

All data needed to evaluate the conclusions in the paper are present in the paper and/or the Supplementary Materials. Additional data related to this paper may be requested from the authors.

## Supplementary Materials and methods

### Energy dispersive X-ray spectroscopy

Samples were prepared for energy dispersive X-ray (EDX) spectroscopy analysis on an aluminium sample holder using adhesive carbon coated tape. Excessive microfibers were removed using pressurized air, after which the samples were sputter coated with 10 nm of carbon. The EDX measurements were performed with a Tescan MAIA III GMH field emission scanning electron microscope (Czech Republic) equipped with a Bruker X-Flash 30 mm^2^ silicon drift energy dispersive X-ray microanalysis detector (MA, USA).

### Micro-Fourier transform infrared spectroscopy

Micro-Fourier transform infrared spectroscopy (μFTIR) measurements were performed using a Thermo Nicolet iN10 micro Fourier transform infrared microscope. Spectra were recorded in the wavelength range from 4000 to 675 cm^−1^ using a spectral resolution of 8 cm^−1^. For the transmission measurements of the polyester reference material and the polyester and nylon environmental fibers, a small amount of the microfibers was transferred onto a diamond micro compression cell where the samples were compressed. For the reflection measurements of the nylon reference material and the polyester and nylon environmental fibers, a small portion of microfibers was suspended in water. The suspension was subsequently filtered over a gold coated 0.8 μm polycarbonate filter (TJ Environmental, The Netherlands). A subset of approximately 100 fibers was individually measured directly on the filter using the reflection mode of the µ-FTIR equipment.

### Extraction of nylon oligomers (mono-, di- and trimer)

A round bottom flask containing 25.1 g cryogenically milled nylon powder (PA66, Sigma-Aldrich) and 500 ml methanol (VWR) was equipped with a reflux condenser and the suspension was stirred overnight at 50°C. Next, the suspension was cooled to approximately 30°C and filtered over a cellulose filter (VWR) to remove the remaining powder. The solvent was removed *in vacuo* using a rotary evaporator (Büchi rotavapor R-215, Switzerland). A white solid was obtained (yield 220 mg), of which the composition was determined using liquid chromatography/mass spectrometry (LC/MS) analysis.

### Isolation of nylon oligomers (mono-, di- and trimer)

A column for silica gel chromatography (⌀ 30 mm, VWR) was charged with silica gel 60 (27 g, 0.063-0.200 mm, Merck) and dichloromethane (DCM, VWR) as eluent. The crude extract containing the mixture of oligomers (200 mg) was added on top of the silica gel column and oligomers were separated on the column using DCM:methanol as eluent (DCM:MeOH gradient: 100:0 → 90:10 → 80:20), which resulted in complete separation of the oligomers. The collected fractions were checked for the presence of product using LC/MS. The fractions containing pure oligomer were combined, filtered over a glass filter (VWR) and subsequently the solvent was removed *in vacuo*. The obtained solids were further dried *in vacuo*, after which the pure oligomers were obtained as white solids. The structure of the oligomers was confirmed by ^1^H NMR spectroscopy.

### Liquid chromatography/mass spectrometry

#### Qualitative analysis of nylon leachate and oligomers

Qualitative analysis of nylon oligomers was performed with an Agilent 1260 series high-performance liquid chromatographer (CA, USA) equipped with a 100 x 2 mm, 3 µm Gemini NX-C18 110 Å LC Column (Phenomenex, Utrecht, The Netherlands), coupled with an Agilent 6410 triple quadrupole LC/MS with electron spray ionization (ESI) in positive SCAN mode. In addition, the LC/MS analysis of the nylon leachate was performed with the Agilent 1260 liquid chromatographer coupled to an Agilent 6460 triple quadrupole LC/MS with Jetstream ESI in positive SCAN mode. A sample volume of 5 µl was injected with a column temperature of 60 °C and a flow rate of 200 µl min^−1^. The sample was eluted with a gradient of Milli-Q water (containing 5 mM ammonium formate with 0.0025% formic acid (both Sigma-Aldrich), eluent A) and methanol (containing 5 mM ammonium formate with 0.0025% formic acid, eluent B) with a flow rate of 0.5 ml min^−1^. Eluent B was increased from 10% to 90% in 10 minutes and maintained for 3 minutes. After this, eluent B was decreased to 10% in 0.1 minute and maintained for 1.9 minute to complete the cycle of 15 minutes. Mass spectrometry was performed with a gas temperature of 350°C and a flow rate of 10 l min^−1^. Stealth gas temperature (for Agilent 6460) was set at 400 °C with a flow rate of 12 l min^−1^. The capillary voltage was set at 4000 V.

#### Direct injection of nylon leachate

An injection volume of 10 µl diluted nylon leachate was directly injected into an Agilent 6410 triple quadrupole MS system with ESI in positive SCAN mode. The conditions were as follows: gas temperature 350 °C, flow rate 10 l min^−1^, mobile phase 50:50 ratio of 80:20 acetonitrile (VWR):Milli-Q water with 5 mM ammonium formate and 10:90 acetonitrile:Milli-Q water with 5 mM ammonium formate, scan range 50–1000 Da, capillary voltage 3500 V.

### ^1^H Nuclear magnetic resonance

The chemical structure of the oligomers was confirmed by ^1^H NMR spectroscopy (Bruker Avance 400 spectrometer). The oligomers were dissolved in ∼0.5 ml CD_3_OD:CDCl_3_ (1:1) (Sigma-Aldrich). The spectra were recorded at 24 °C, and internally referenced to the residual solvent resonance (CD_3_OD: ^1^H *δ* 3.31).

<u>Nylon monomer</u>, yield = 77 mg. ^1^H NMR (400.1 MHz, CD_3_OD/CDCl_3_ 50:50, 297 K) δ = 7.56 (br. m, 2H; N*H*CO), 3.22 (m, 4H, NHC*H*_2_), 2.19 (m, 4H, COC*H*_2_), 1.63 (m, 4H, COCH_2_C*H*_2_), 1.54 (m, 4H, NHCH_2_C*H*_2_), 1.32 (m, 4H, NH (CH_2_)_2_C*H*_2_).

<u>Nylon dimer</u>, yield = 74 mg. ^1^H NMR (400.1 MHz, CD_3_OD/CDCl_3_ 50:50, 297 K) δ = 7.68 (br. m, 4H; N*H*CO), 3.15 (t, 8H, NHC*H*_2_), 2.18 (m, 8H, COC*H*_2_), 1.60 (m, 8H, COCH_2_C*H*_2_), 1.47 (m, 8H, NHCH_2_C*H*_2_), 1.31 (m, 8H, NH (CH_2_)_2_C*H*_2_).

<u>Nylon trimer</u>, yield = 16 mg. ^1^H NMR (400.1 MHz, CD_3_OD/CDCl_3_ 50:50, 297 K) δ = 7.74 (br. m, 6H; N*H*CO), 3.14 (t, 12H, NHC*H*_2_), 2.18 (m, 12H, COC*H*_2_), 1.60 (m, 12H, COCH_2_C*H*_2_), 1.48 (m, 12H, NHCH_2_C*H*_2_), 1.32 (m, 12H, NH (CH_2_)_2_C*H*_2_).

## Supplementary tables and figures

**Table S1.**
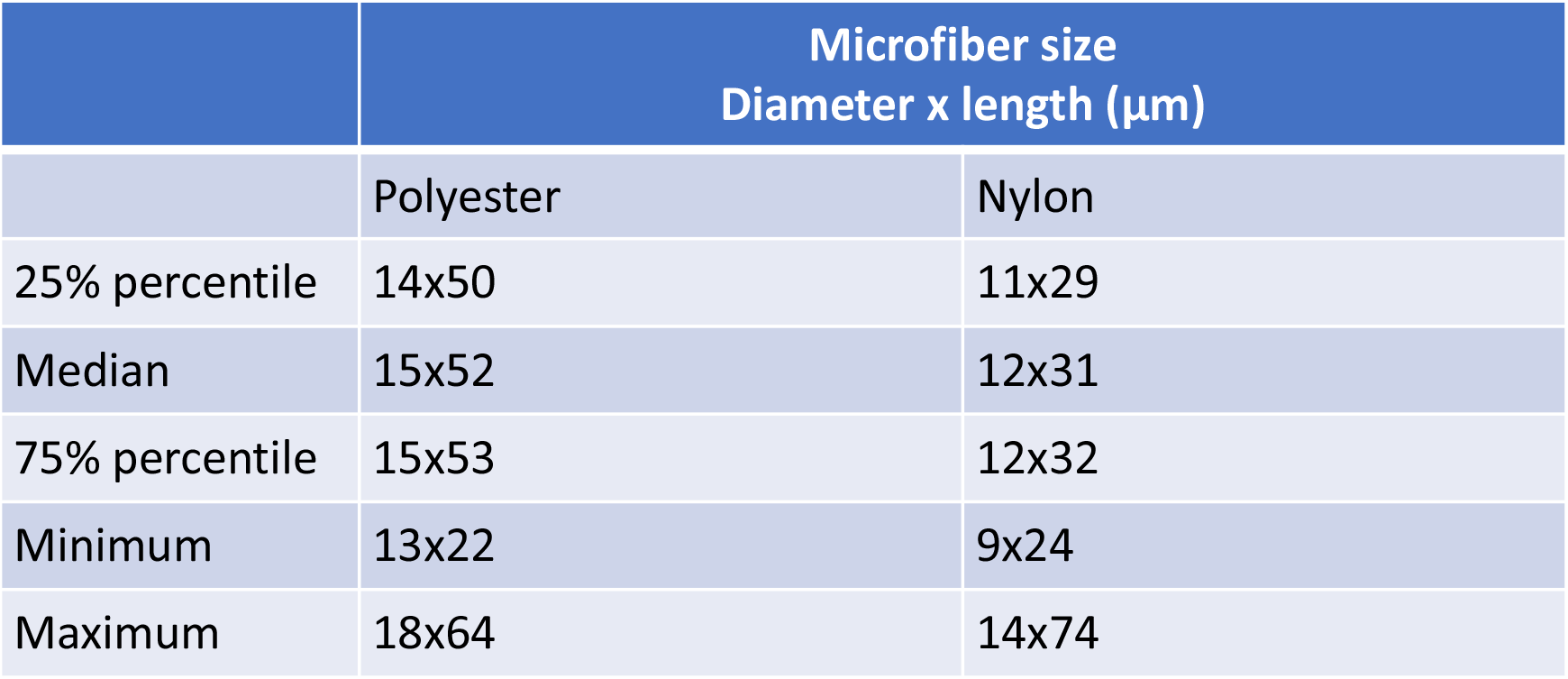
Size characteristics of polyester and nylon reference microfibers.

**Figure S1:**
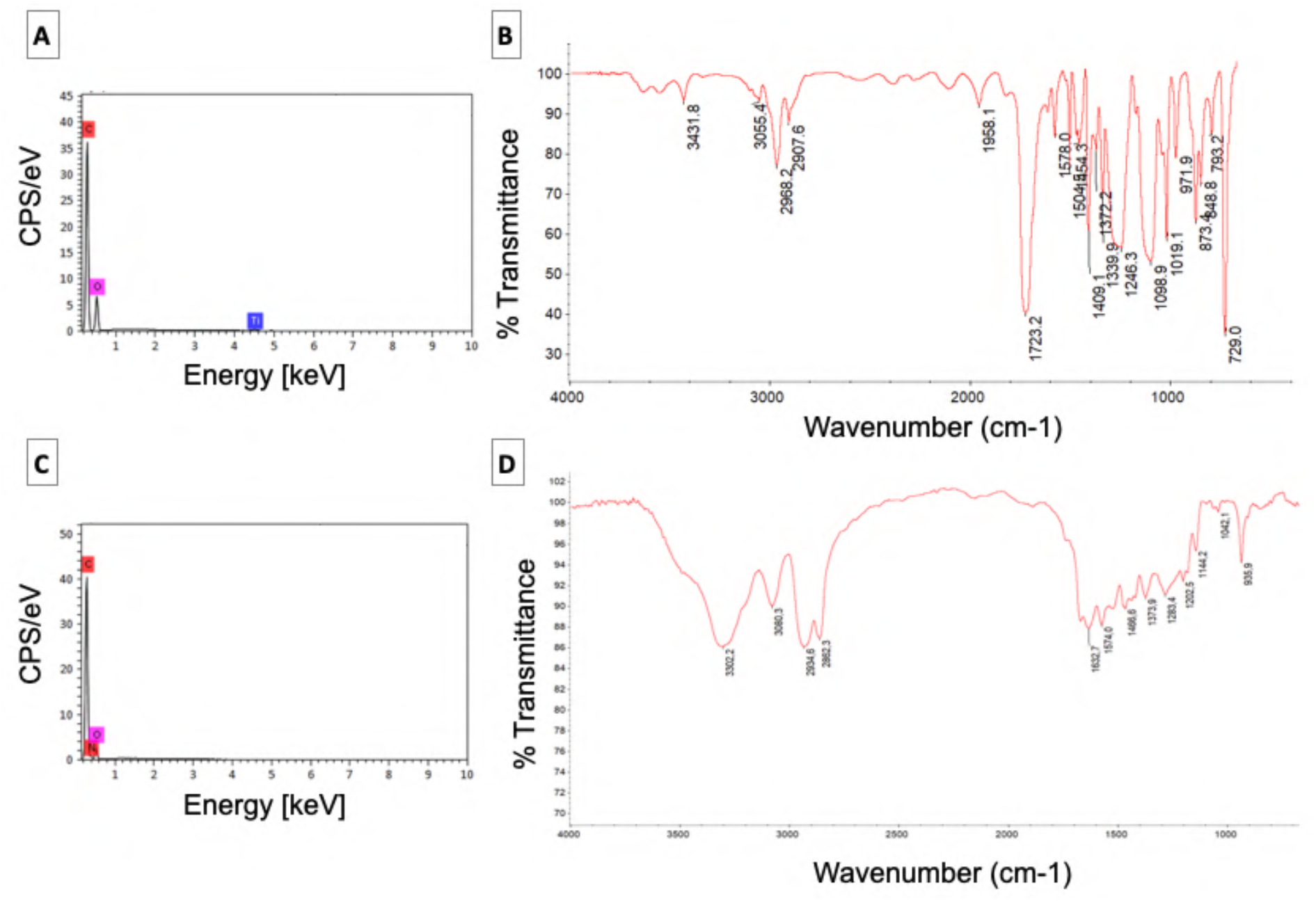
Characterization of the reference microfibers using energy dispersive X-ray - and infrared spectroscopy. (**A**) EDX spectrum of polyester, confirming the presence of carbon (C) and oxygen (O), and additionally revealing the presence of titanium (Ti), which can be ascribed to small TiO2 pigment particles used as filler material in these fibers. (**B**) µFTIR spectrum of polyester with characteristic absorbance peaks (2968 cm^−1^, C-H stretch; 1723 cm^−1^, C=O stretch; 1246 cm^−1^, C-O stretch aromatic ester; 729 cm^−1^, benzene derivative (79)). (**C**) EDX spectrum of nylon, confirming the presence of carbon (C), nitrogen (N) and oxygen (O). (**D**) µFTIR spectrum of nylon with characteristic nylon absorbance peaks (3302 cm^−1^, N-H stretch; 2934 cm^−1^, C-H stretch; 1632 cm^−1^, C=O stretch sec. amide; 1202 cm^−1^, C-N bend (79)).

**Figure S2:**
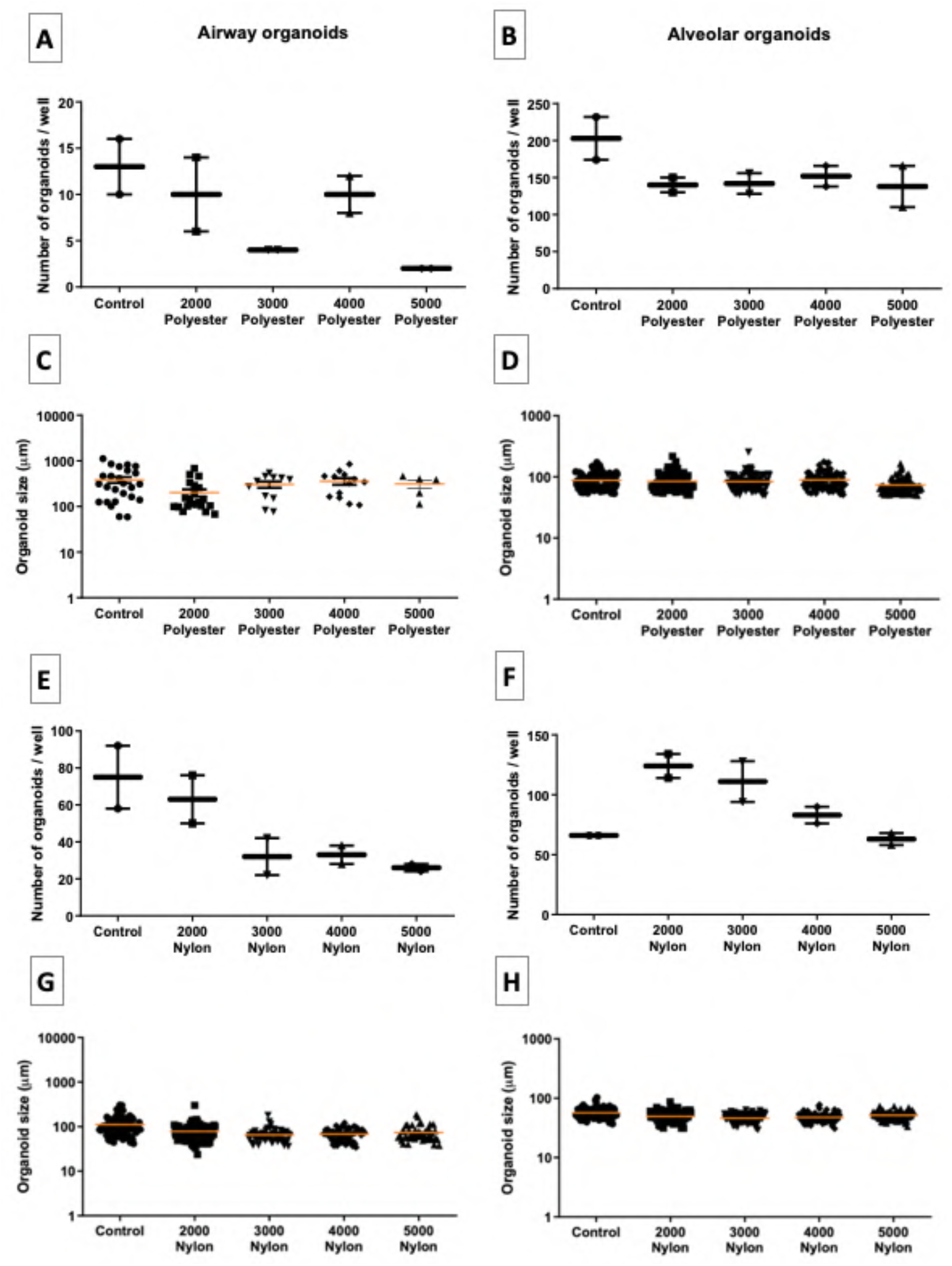
Determination of the optimal microfiber dose for subsequent in vitro testing of polyester and nylon reference microfibers, using 2000, 3000, 4000 and 5000 microfibers per condition. **(A, E)** Quantification of airway and **(B, F)** alveolar organoid numbers for organoids exposed to polyester or nylon, respectively (n=2 independent isolations). **(C, G)** Quantification of the airway and **(D, H)** alveolar organoid size following exposure to polyester or nylon. 2000, 3000, 4000 or 5000 fibers per well corresponded to approximately 49, 73, 98 and 122 μg/ml of polyester fibers or 16, 23, 31, and 39 μg/ml of nylon fibers.

**Table S2.** Size characteristics of polyester and nylon environmental microfibers.

| | Microfiber size<br>Diameter x length ($\mu\text{m}$) | |
| --- | --- | --- |
|  | Environmental polyester | Environmental nylon |
| 25% percentile | 15x54 | 46x15 |
| Median | 17x63 | 57x20 |
| 75% percentile | 18x85 | 73x27 |
| Minimum | 8x30 | 17x8 |
| Maximum | 24x269 | 296x66 |

**Figure S3:**
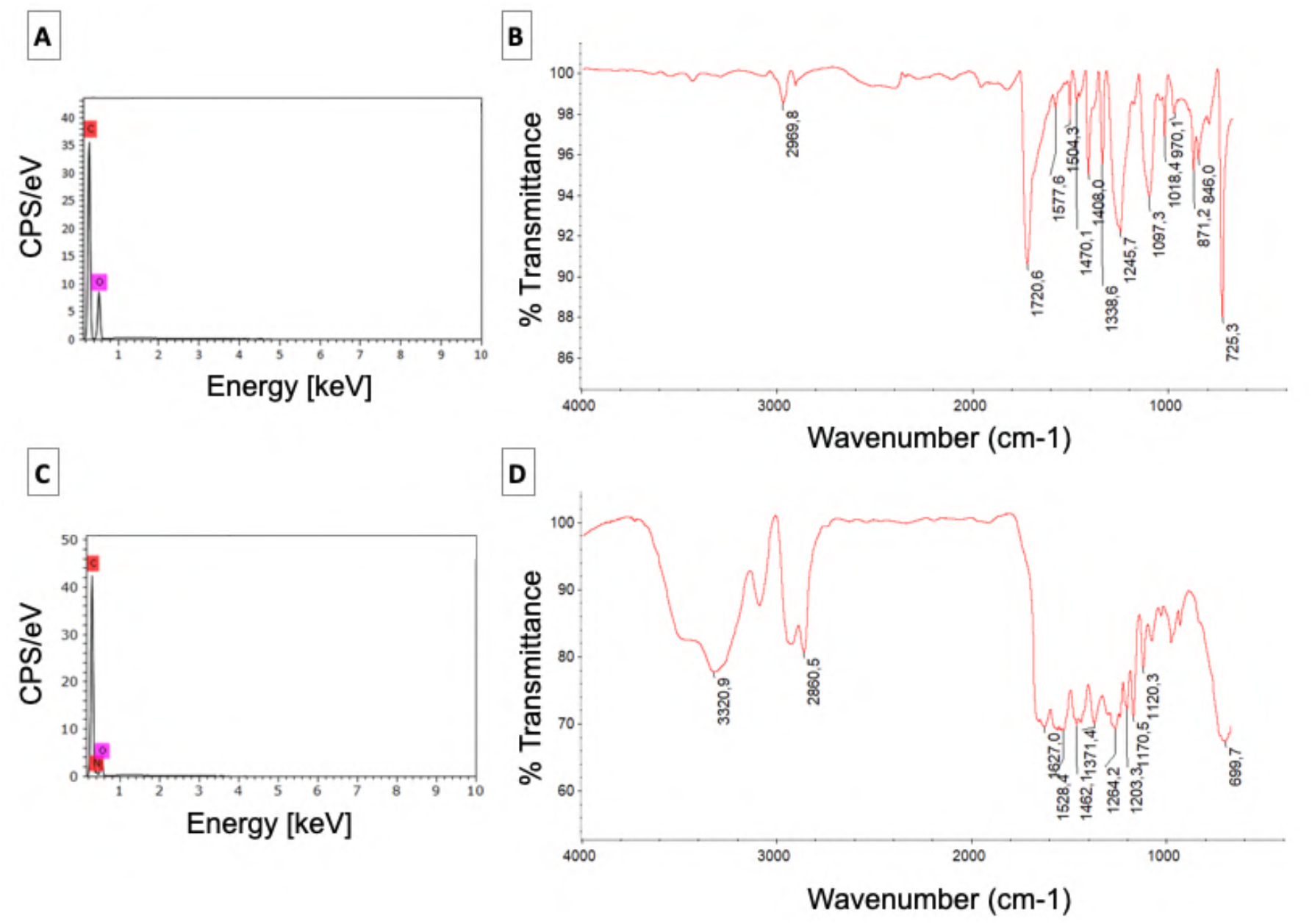
Characterization of the environmental microfibers using energy dispersive X-ray - and infrared spectroscopy. EDX - and µ-FTIR spectra of (**A** and **B**) polyester and (**C** and **D**) nylon microfibers.

**Figure S4:**
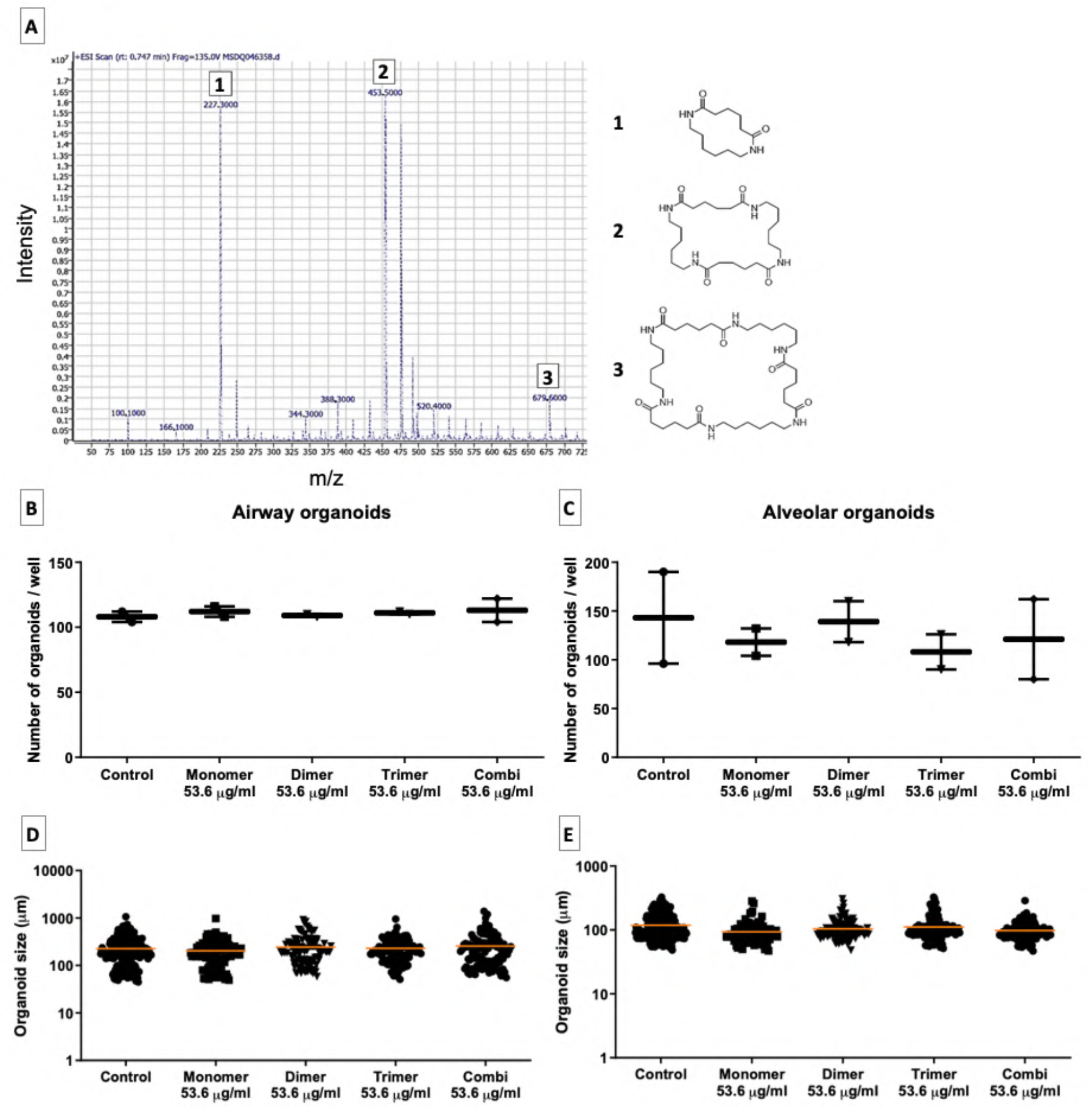
Characterization of the components leaching from nylon reference microfibers and their effect on organoid growth. (**A**) Mass spectrometry spectrum of the nylon leachate, revealing high amounts of cyclic nylon mono-, di- and trimers, as well as other smaller peaks. (**B** and **C**) Assessment of the numbers of airway and alveolar organoids and (**D** and **E**) their sizes (n=2 independent isolations).

**Figure S5:**
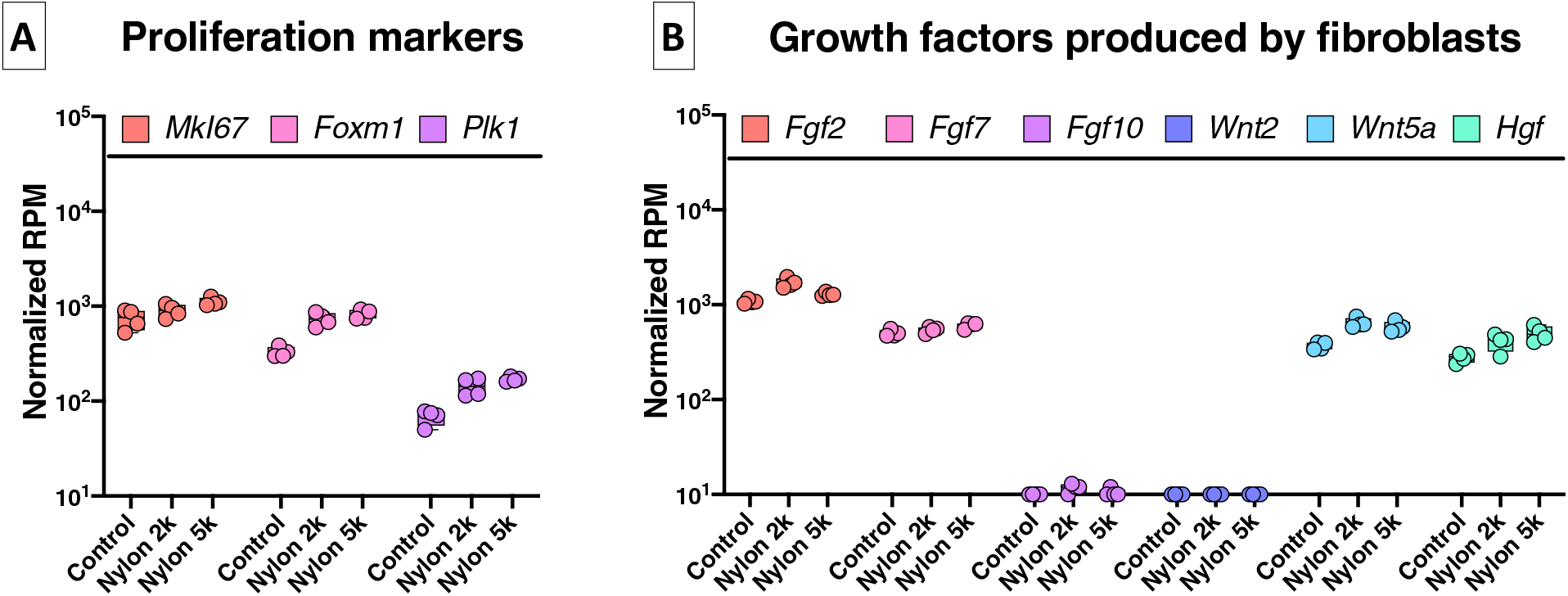
Expression of individual genes in fibroblasts isolated from organoid cultures exposed to nylon microfibers. (**A**) Genes associated with proliferation of fibroblasts. (**B**) Genes encoding factors produced by fibroblasts important for epithelial development (n=4 independent isolations). 2k= 2000 fibers, 5k=5000 fibers

